# Cell4D: A general purpose spatial stochastic simulator for cellular pathways

**DOI:** 10.1101/2023.09.10.557076

**Authors:** Donny Chan, Graham L. Cromar, Billy Taj, John Parkinson

## Abstract

**Motivation:** With the generation of vast compendia of ‘omics datasets, the challenge is how best to interpret these datasets to gain meaningful biological insights. Key to this challenge are computational methods that enable domain-users to generate novel hypotheses that can be used to guide future experiments. Of particular interest are flexible modeling platforms, capable of simulating a diverse range of biological systems with low barriers of adoption to those with limited computational expertise.

**Results:** We introduce Cell4D, a spatial-temporal modeling platform combining a robust simulation engine with integrated graphics visualization, a model design editor, and an underlying XML data model capable of capturing a variety of cellular functions. Cell4D provides an interactive visualization mode, allowing intuitive feedback on model behaviour and exploration of novel hypotheses, together with a non-graphics mode, compatible with high performance cloud compute solutions, to facilitate generation of statistical data. To demonstrate the flexibility and effectiveness of Cell4D, we investigate the dynamics of CEACAM1 localization in T-cell activation. We confirm the importance of Ca^++^ microdomains in activating calmodulin and highlight a key role of activated calmodulin on the surface expression of CEACAM1. We further show how lymphocyte-specific protein tyrosine kinase can help regulate this cell surface expression and exploit spatial modeling features of Cell4D to test the hypothesis that lipid rafts regulate clustering of CEACAM1 to promote trans-binding to neighbouring cells. Through demonstrating its ability to test and generate hypotheses, Cell4D represents an effective tool to help interpret complex ‘omics datasets.

**Availability and Implementation:** https://github.com/ParkinsonLab/cell4d

## INTRODUCTION

The post-genome era is characterized by increasing availability of large, heterogeneous datasets detailing the molecules driving biological systems. These include genome scale datasets encompassing the expression and dynamics of genes and their products [1-3], their localization [4, 5], interactions with other biomolecules [6, 7] and organization within pathways [8, 9]. A major challenge is how best to unlock the full potential of these rich datasets to advance our understanding of complex biological processes. To address this challenge, a variety of computational modeling platforms have been developed, capable of exploiting these datasets to simulate the dynamics of the underlying systems at the meso-scale [10-12] [13].

Despite the availability of these platforms, widespread adoption has been limited, in part due to barriers concerning the level of computational expertise required for their operation. To help overcome these limitations, toolkits such as CellBlender [14], have been developed to help with the construction and visualization of models. At the same time there is a need for simulation platforms for cell biologists, without skills in computing, who nevertheless represent the domain experts and target audience for meso-scale simulators. To meet this need, we have developed Cell4D, a robust spatial stochastic simulation platform with integrated graphical visualization and a browser-based editing tool supporting the creation of sophisticated models written in human-readable XML format, ensuring cross-platform compatibility and allowing for further feature enhancements.

To demonstrate the ability of Cell4D to model a biological system and explore hypotheses, we applied it to examine the dynamics of carcinoembryonic antigen-related cell adhesion molecule 1 (CEACAM1) in T-cell activation [15]. CEACAM proteins are glycosylated transmembrane adhesion molecules featuring an N-terminal IgV-like domain together with a variable number of IgC2-like domains, a transmembrane domain and a cytoplasmic tail [16-18]. They are widely expressed in many cell types and a play a role in multiple functions including cell growth, metabolism and responding to infection. CEACAM1 is expressed by activated T cells and serves to transmit extracellular signals across the cell membrane through intercellular (trans) homophilic and heterophilic binding through its transmembrane domain to inhibit continued activation [19]. In resting T cells, CEACAM1 largely exists in the form of cis-homodimers, incapable of binding extracellular ligands. Clathrin-mediated internalization further ensures a relatively low concentration at the cell surface. Upon binding by active calmodulin, cis-homodimers disassociate facilitating trans-binding of the monomeric form of CEACAM1 to other monomers of CEACAM1 on adjacent cells [19]. Such binding appears higher in the presence of multiple IgC2 domains, suggesting a role for local accumulation (clustering) of CEACAM1 monomers [16, 20]. Regulation of CEACAM functionality is thought to operate through localized concentrations of Ca^2+^ [21] as well as phosphorylation of CEACAM1 by Src-family kinases such as Lck [22]. Furthermore, it has been hypothesized that regulation may also involve the preferential sequestration of CEACAM1 monomers within membrane microdomains [23].

Applying Cell4D we examine the dynamics of calmodulin activation through calcium binding and subsequently integrated this model to examine the role of calmodulin, Lck kinase and membrane microdomains to regulate the formation of local surface clusters of monomeric CEACAM1 to promote trans-binding.

## SYSTEM AND METHODS

Cell4D is written in C++. To perform simulations, model files are loaded into the simulation environment which features a graphical interface that allows for real-time visualization of the running simulation. To help with the generation of model files, a Cell4D Model Editor - accessible at https://compsysbio.org/cell4d_ui/build/ is provided. Simulations iterate over a user-defined number of timesteps. Within each timestep, molecular diffusion events are first simulated followed by reaction events (**Figure 1**). For small molecules represented by local concentrations within a lattice cell (c-voxel), the diffusion into neighboring c-voxels are calculated deterministically [24]. For molecules and complexes represented as point particles, displacement is determined either stochastically (i.e., Brownian motion) or deterministically (i.e. active transport). Full Details of the System and Methods are provided in **Supplemental Text**.

**Figure 1:**
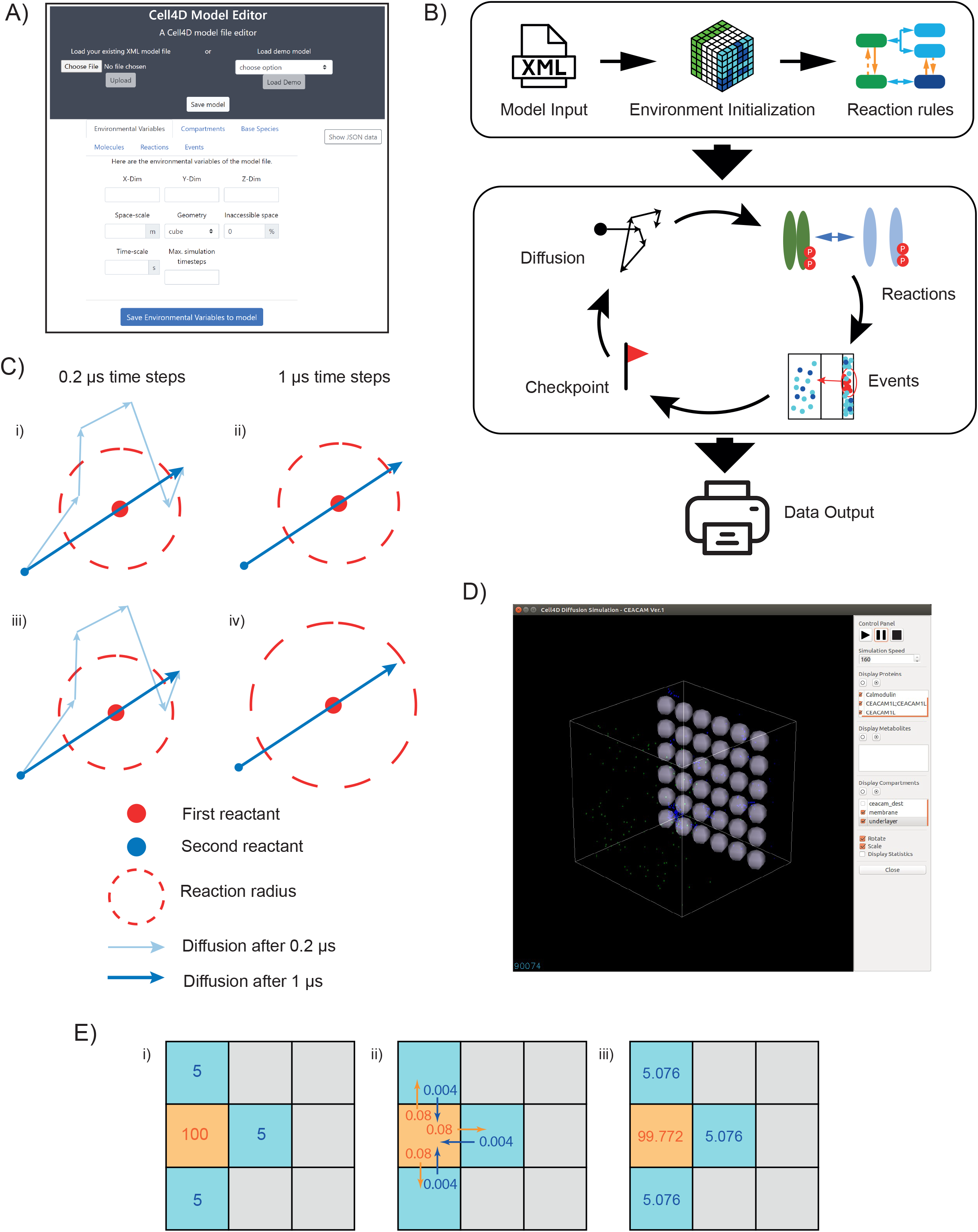
Conceptual overview of Cell4D. A) Model design interface. A web interface for creating and editing Cell4D model files. Custom XML model files can be loaded in by a user, or a preset example model can be selected. B) Simulation set up and flow time cycle logic. Parameters that describe system behavior such as the way molecules behave within the simulation space as well as interact with other molecules are described in an XML input file, which is then used to initialize the simulation space. The simulation then cycles through a series of steps until the end condition is met. Output occurs in two forms: tab-delimited files (.tsv) of molecule counts data at each time step, while particle logs record the position and state information of molecules in the simulation. C) Implementation of off-lattice movement of point particles. While short time steps (here 0.2µs) allow for reactions between two particles within a specified reaction radius, longer time steps (here 1µs) may result in missed reactions. To avoid such cases Cell4D implements the Andrews-Bray-adjustment to artificially increase reaction radii of molecules [26]. D) Screenshot of the Cell4D graphical interface. E) Diffusion of bulk molecules shown for a single voxel. At each time step, a portion of the bulk molecules for each c-voxel will diffuse into a neighboring c-voxel, based on the current concentration and the molecule’s diffusion rate constant. This is calculated for all c-voxels at every timestep.

## ALGORITHMS

### Molecular movement and reactions

Point particle movement is modeled through random walks, while bulk particle movement is modeled using Fick’s laws of diffusion [24] and is simulated using the forward Euler method [25]. Reactions are implicitly defined based on presence of substrates. Cell4D allows the definition of unimolecular and bimolecular reactions. For bimolecular reactions involving point particles, Cell4D uses the Andrews-Bray-adjusted Smoluchowski method. For further details see **Supplemental Text**.

### Modeled Systems

#### Calmodulin Activation

Models used a simulation environment defined as a cube with side length of 8 × 10^−7^ m. Calcium ions were defined as bulk molecules and calmodulin was defined as point particles with N- and C-terminal bindings sites. For some simulations, The model space was divided into a lattice comprising 10 × 10 × 10 c-voxels which were subdivided into five 2 × 10 × 10 cross-sectional compartments (quintiles Q1 thru Q5). Within Q1, for c-voxels were defined as sites of calcium-release.

#### CEACAM1 activation

Models used a simulation environment composed of 6 × 6 × 6 c-voxels of length 0.2 μm, subdivided into an outer ‘membrane’, a ‘cytosolic interface’, a ‘cytosol’ and an ‘organelle’. For some simulations, within the membrane, microdomains were defined to represent lipid rafts. The model features CEACAM1 molecules capable of forming monomers or dimers, Lck molecules capable of phosphorylating CEACAM1, calmodulin which can dissociate CEACAM1 dimers to monomers and calcium, which can activate calmodulin.

Details of both systems are provided in **Supplemental Text**.

## IMPLEMENTATION

### Cell4D accurately simulates diffusion and reaction events

Cell4D is a spatio-temporal simulation platform that performs simulations within a space defined by cubic lattice sites (c-voxels). The simulator features reaction-based rules governing molecular interactions, formation and dissolution of protein complexes, state changes (e.g., allowing post-translational modifications), enzymatic reactions and defined events (e.g., molecule trafficking between compartments). Using a hybrid on/off lattice approach, small molecules are represented as concentrations, diffusing via a grid pattern dividing the overall model space while larger molecules can be tracked as point particles that freely diffuse off-lattice. Within the lattice, multiple compartments can be defined (e.g., membranes or other organelles), providing boundaries and allowing the definition of compartment-specific rules governing molecule behavior. Compartments can contain other compartments and both compartments and the lattice environment itself can be trimmed to allow the representation of custom geometries. All these rules may be combined within the same data model without the need for custom modifications to the code.

In initial simulations we benchmarked the ability of Cell4D to accurately model molecular diffusion and reaction events (see **Supplemental Text**). Focusing on diffusion, we found that Cell4D accurately models Brownian motion for both point particles and bulk molecules as predicted by Fick’s laws (**Supplemental Figures 1 & 2**). In terms of reactions, simulated product formation for unimolecular reactions matched theoretical yields under all tested conditions (**Supplemental Figure 3**). For bimolecular reactions involving both bulk molecules and point particles, we found that while timescale had negligible impact, accuracy increased for simulations with higher rates of diffusion (**Supplemental Figure 4**). Bimolecular reactions involving only point particles were found to be sensitive to timescale and reaction rate (**Supplemental Figure 3B**). To correct these errors we implemented the Andrews-Bray (AB) adjustment of the Smoluchowski method (**Supplemental Figure 5**), resulting in high accuracy predictions under all conditions except those violating the assumption that the main constraint for reaction rates stems from particle collision [26]. Having benchmarked the performance of Cell4D, we next demonstrate the ability of Cell4D to simulate a biological system.

### Application of Cell4D to investigate mechanisms underlying CEACAM1 mediated T-cell activation

We chose to focus on CEACAM1 mediated T-cell activation to examine two aspects of Cell4D’s capacity to model biochemical pathways. First, we use Cell4D to explore the dynamics of calcium:calmodulin binding dynamics, a critical step in the activation of CEACAM1 signaling.. Second, we use Cell4D to examine the hypothesis that CEACAM1 signaling is regulated through its spatial organization at the cell surface and the underlying mechanisms that compete to determine its local concentration.

### Calcium microdomains play a key role in calmodulin activation

To model calmodulin activation we applied a cooperative binding model [27] to confirm the importance of calcium microdomains in overcoming cellular buffering in signal transduction. Details on the cooperative binding kinetics involving the four calcium binding domains of calmodulin are provided in **Supplemental Text**. First we examined a baseline model to establish that calmodulin binding occurs under physiological calcium concentrations. To compare simulations with theoretical predictions, we initially predicted the distribution of the four potential states of calmodulin: CaM_1, CaM_2, CaM_3, and CaM_4, representing calmodulin molecules with 1, 2, 3, or 4 binding sites occupied with calcium respectively, for 2µM calmodulin exposed to a range of Ca^2+^ concentrations (0 to 16μM) in a volume of 5 × 10^−16^ L. Calculations were performed using the forward Euler method under the well-mixed assumption (see **Supplemental Text**). The same conditions were then reproduced using Cell4D by placing the same concentration of molecules distributed randomly in a simulation space of 4 × 4 × 4 c-voxels of length 0.2 μm. Results from simulations were similar to theoretical predictions and showed that under Ca^2+^ concentrations relevant to T cell activation (100nM – 1.2µM), the proportion of saturated (activated) calmodulin (CaM_4) was negligible (<1%; **Supplemental Figure 6**). Instead, we found that modest activation of calmodulin (~20%) required Ca^2+^ concentrations an order of magnitude higher (>10µM) than observed experimentally [28, 29].

While the average concentration of calcium in a cell is approximately 1-2 µM, local concentrations can be significantly higher (of the order 100µM – 1mM) in regions proximal to active calcium channels, which we term *microdomains* [27]. To explore the range and impact of such channels on calmodulin activation, we constructed a model composed of five compartments (defined as quintiles, Q1-Q5) each composed of 2 × 10 × 10 c-voxels of 0.08 µm. 1mM of Ca^2+^ was then introduced in Q1 at each timestep and allowed to freely diffuse and exit from Q5 (**Figure 2A**). To assess the potential impact of the spatial organization of Ca^2+^ release, three arrangements were investigated (see **Supplemental Text**). As expected, Ca^2+^ concentrations decreased with increasing distance from the source, ranging from 1-10µM across the five quintiles (**Figure 2B**). Further, the relative proportion of activated calmodulin (CaM_4) reflects the distance from the source of Ca^2+^ (from 5-10% and 20-30% for Q5 and Q1 respectively; **Figure 2C**). Configuration of calcium channels had minimal impact on either calcium concentrations or activated calmodulin. Interestingly, in Q5, despite the proportion of primed calmodulin (CaM_2) being ~75%, activated calmodulin remained relatively low (5-10%). This suggests that while elevated background concentrations of Ca^2+^ are sufficient to maintain a primed population of calmodulin, its activation appears transient and likely requires proximity to sources of Ca^2+^, a local effect that is not well captured in well-mixed models.

**Figure 2:**
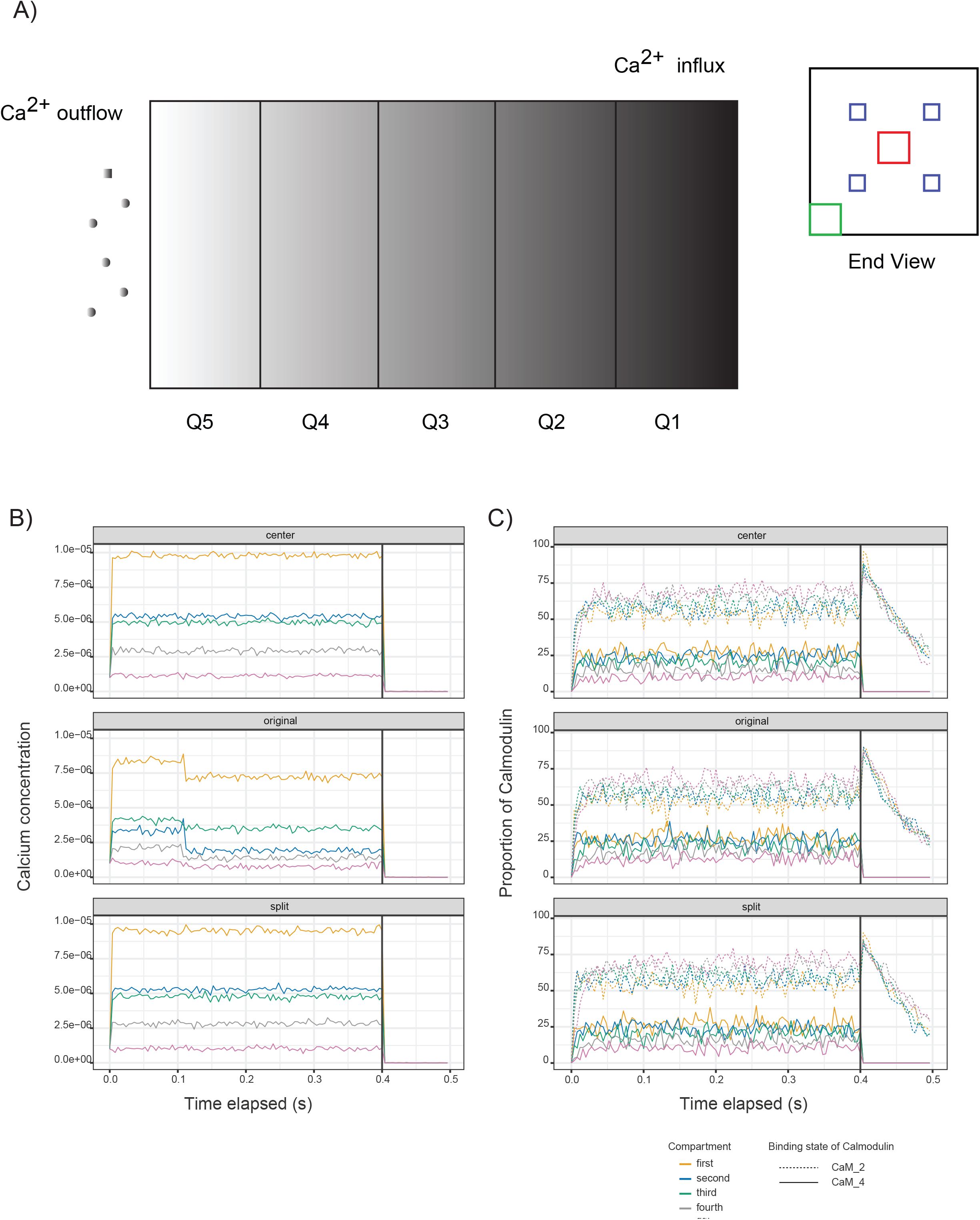
Activation of calmodulin in Ca^2+^ microdomains. A) Schematic representation of simulations of calmodulin activation involving five compartments (Q1 – Q5). End view shows the spatial orientation of the three types (indicated by green, red and blue squares) of microdomain-layout used to release Ca^2+^ into the system. Calcium is removed at the Q5 region to maintain the expected overall Ca^2+^ concentration of approximately 1-2 µM. B) Effective average concentration of Ca^2+^ in each of the compartments over the course of the simulations. C) Average saturation of two states of calmodulin (CaM_2 and CaM_4) in each compartment over the course of the simulations.

Having established the conditions under which elevated Ca2+ levels lead to calmodulin activation, we next consider factors affecting the spatial organization of CEACAM1 in response to T-cell activation.

### Clustering of CEACAM1 at the cell surface indicates a role for lipid rafts in regulating signalling and is dependent on calmodulin activation

The primary mechanism of CEACAM1 function is to transduce extracellular signals to the cytosol through intracellular *trans*-binding. In resting T cells, CEACAM1 concentration at the cell membrane remains relatively low due to clathrin-mediated internalization, involving the adaptor proteins AP1 and AP2. CEACAM1 in resting T cells is primarily found in the cis-dimeric form, predicted to be inactive due to steric hinderance that prevents access to the ITIM sites in the cytoplasmic tail, blocking SHP1 recruitment which leads to downstream CEACAM1 inhibitory function (not considered here). Upon binding with calmodulin, CEACAM1 dimers are converted to their monomeric form. Phosphorylation of tyrosine residues in the ITIM sites by Lck, both blocks binding to AP1 and AP2 adaptor proteins (preventing internalization and resulting in retention of CEACAM1 at the cell surface) as well as promote binding to their counterparts on adjacent cells in *trans* (resulting in the suppression of host immune responses). With the observation of CEACAM1 clustering at the cell surface, it has further been suggested that the CEACAM1 binding dynamics may also be driven, at least in part, by partitioning through lipid rafts [16, 21, 23]. Here we are interested in using Cell4D to examine potential mechanisms, including interactions involving calcium, calmodulin, Lck kinase as well as the involvement of lipid rafts, that drive the accumulation of CEACAM1 at the cell surface (**Figure 3A**).

**Figure 3:**
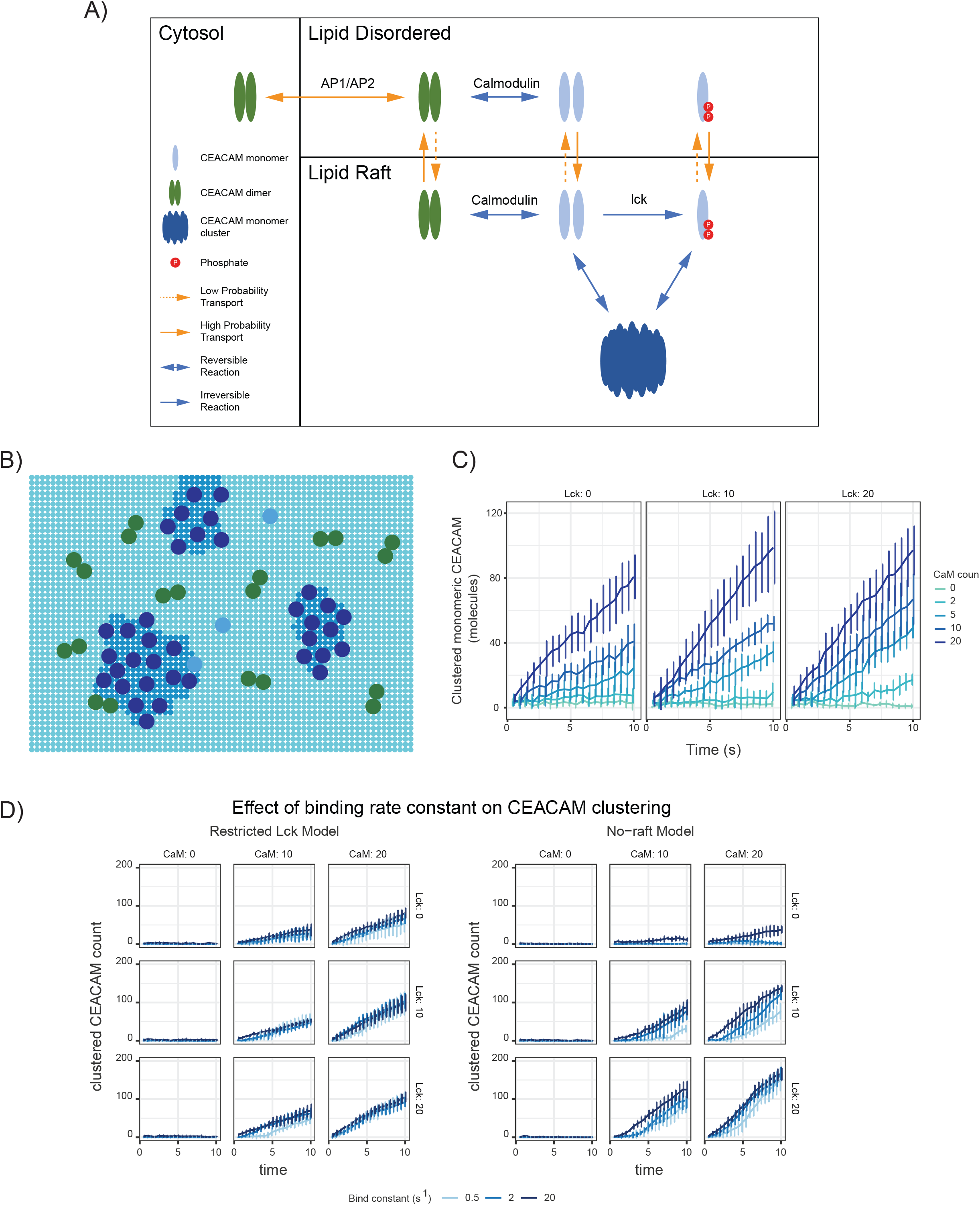
Modeling CEACAM1 signaling. A) Schematic of reactions used in the model. CEACAM1 dimers are transported between the membrane and cytosol compartments. Within the membrane, CEACAM1 dimers disassociate to monomers based on interactions with activated calmodulin. Src-family kinases (Lck) phosphorylate the ITIM regions of the CEACAM1 cytoplasmic tail, preventing its transport back to the cytosol and shifting the equilibrium of CEACAM1 localization to the membrane. The membrane region can be defined into lipid-ordered (lipid rafts) and lipid-disordered regions. CEACAM1 preferentially associates with these regions based on its oligomeric state, as shown by the solid and dashed orange arrows between the two membrane regions that indicate transport. The end state of the activated T cell consists of the clustering of CEACAM1 monomers within lipid rafts. B) Representation of 2D membrane compartment with lipid raft sub-compartments. C) The lipid raft CEACAM1 model was tested using 0, 10, 20 molecules of Lck and 0, 2, 5, 10, 20 molecules of active calmodulin to examine the effects of both proteins on CEACAM1 surface expression. Error bars represent standard deviation for 6 replicates. Results show that CEACAM1 surface concentration is dependent on the concentration of activated calmodulin, but not Lck. D) Impact of trans-binding rate constants (i.e. unbound monomer to trans-bound (immobilized) monomer) on CEACAM clustering. Simulations were performed for both the lipid raft model (left) and the no-raft model (right). The binding rate constant shows a positive correlation with the total count of trans-bound (clustered) CEACAM1 monomers. For low binding constant conditions in the presence of Lck and CaM in the no-raft model, there appears to be a threshold effect where the rate of cluster formation is slow in the beginning of the simulation, but accelerates after a certain point. For the no-raft Lck-absent models, low calmodulin levels did not lead to a significant clustered CEACAM1 population, while high calmodulin only produced an increased surface CEACAM1 concentration at the highest binding rate constant.

To examine the impact of lipid rafts, we constructed two models (*raft* and *no-raft* models). Both models involve an environment composed of 6 × 6 × 6 voxels of length 0.2μm sub-divided into four compartments representing the cell membrane, a region of cytosol immediately adjacent to the membrane, the rest of the cytosol and an organelle (see **Supplemental Text**). Particles in the membrane were restricted to diffusion in 2-dimensions only. Calmodulin was confined to the cytosolic interface with the membrane while particles representing CEACAM1 could move between compartments through defined transport events (representing clathrin-mediated internalization and export to the membrane). In the raft model, CEACAM1 monomers are preferentially enriched in membrane microdomains, while CEACAM1 dimers are preferentially excluded. Further, Lck kinases, previously shown to be enriched and functionally segregated in lipid rafts [30], were constrained within microdomains, representing rafts, defined within the membrane (**Figure 3B**). Thus, in this model, Lck is only able to phosphorylate CEACAM1 (through its ITIM domain) located within the lipid rafts. Simulations were performed for 200,000 timesteps of 50 μs (10 seconds total) and explored the impact of different concentrations of Lck and Ca^2+^-activated calmodulin, on the concentration and distribution of CEACAM1. Models initiated with no calmodulin or Lck represent the T cell model at rest, while the T cell activated state was modeled through the inclusion of 20 molecules of activated calmodulin and Lck (0.1μM).

For both raft and no-raft models, surface clustering of CEACAM1, representing *trans* bound CEACAM1, was found to depend on the availability of activated calmodulin (CaM_4; **Figure 3C**). However, counter-intuitively, in the absence of rafts, the total amount of CEACAM1 localized to the membrane exceeded that of the raft model (**Supplemental Figure 7**). One possible explanation is that the preferential accumulation of monomeric CEACAM1 in the rafts may increase the relative concentration and promote self-association into homodimers, thus reducing the amount of monomeric CEACAM1 available for phosphorylation by Lck. Where such unexpected behaviors emerge, there is an opportunity to question the model and its assumptions. Here, possible refinements could include changes in the relative rates of CEACAM transport, trans-binding or, the incorporation of additional components modeling the activation and partitioning of Lck into lipid rafts.

To further explore this behavior, we examined the impact of CEACAM1 dimerization rate constants in both the raft and no raft models, on CEACAM1 clustering (**Figure 3D)**. For most simulations, the number of clustered CEACAM1 molecules (defined as monomers of CEACAM1 that have been transiently and reversibly immobilized on the membrane -mimicking trans-binding) increased over the course of the simulation. Unlike the no raft model in which clustering of CEACAM1 was sensitive to the dimerization rate constants, such constants had minimal impact in the raft model. This suggests that for CEACAM1 signaling, in addition to regulating local concentrations of Lck, lipid rafts may also play an important role in modulating the ability of CEACAM1 to form clusters, reducing its sensitivity to reaction kinetics.

## DISCUSSION

We present Cell4D, a novel tool for the spatiotemporal simulation of biological processes and pathways. Combining deterministic, cellular automata for the efficient diffusion of small molecules (as concentrations within a virtual lattice), with probabilistic Brownian motion of discrete molecules, Cell4D is a feature-rich and flexible program that can accurately simulate a variety of biological pathways. In our initial simulations, we validated the ability of Cell4D to accurately model molecular diffusion and reactions, the latter exploiting the Andrews-Bray (AB) adjustment of the Smoluchowski method [26]. Next, we applied Cell4D to model two complementary aspects of CEACAM1-mediated signalling: a calcium-calmodulin interaction model and a CEACAM1 localization model. Previously it has been suggested that monomeric CEACAM1 clusters at the cell surface to help amplify CEACAM1-mediated signalling, potentially though the formation of a lattice-like arrangement of trans-dimers involving CEACAM1 monomers on neighbouring cells[16, 21, 23]. Our simulations support the requirement for local and transient spikes in [Ca^2+^] to activate primed calmodulin, and subsequently predict a dependence of CEACAM1 cluster size and surface concentration on active calmodulin. Furthermore, we showed that competing mechanisms have the potential to influence CEACAM1 clustering including the sequestration of Lck and CEACAM1 within lipid rafts, which at the same time may require a threshold amount of CEACAM1 to maintain the formation of clusters. In addition to representing an important biological process, with the involvement of molecular diffusion, spatial compartmentalization, active transport, reactions and state changes, CEACAM1 signaling represents suitably complex system to test the functional capabilities of Cell4D.

It has been previously shown that CEACAM1 clusters in lipid raft regions are predominantly monomeric, while CEACAM1 outside of lipid rafts primarily exists in a dimeric state [31]. To replicate this behaviour in Cell4D, we defined microdomains, representing lipid rafts, to preferentially accumulate monomeric CEACAM1 (and Lck) and exclude dimeric CEACAM1. Furthermore, in our model CEACAM1 monomers have a low probability of spontaneously becoming immobilized (representing a trans-binding event). This combination of accumulation and immobilization amplifies the localization of monomers within lipid-rafts. Furthermore, through increasing the local concentration of CEACAM1 monomers, clustering is promoted.

Thus, the presence of lipid raft regions in the Cell4D models creates a spatial dynamic that matches the known observation of CEACAM1 clusters being primarily monomeric within lipid rafts, and dimeric CEACAM1 being localized in lipid-disordered regions outside of rafts [32].

We acknowledge that our models are based on hypothesized mechanisms as well as being subject to inevitable modeling constraints. For example, since calcium ions diffuse quickly and interact with calmodulin at fast kinetic rates (10^7^ to 10^10^ M^−1^s^−1^), the system resolves quickly requiring less simulation time. However, these kinetic rates necessitate a high-resolution time scale to avoid missing reactions between timesteps. In comparison, the localization model involving CEACAM1 transport, with molecular species that have slower diffusion rates and slower reaction rates (10^4^ to 10^7^ M^−1^s^−1^), require more simulation time to reach equilibrium. The solution was the creation of separate, complimentary models, which apart from demonstrating Cell4D’s modeling flexibility, allowed each part of the pathway to be simulated with appropriate simulation time and space scales.

We further note that the simulation of vesicle formation and transport of CEACAM1 between compartments was simplified using a single, probabilistic transport event. Future modifications of Cell4D are anticipated to support dynamic compartments (i.e., compartments which move, merge with, or bud from other compartments). In initial experiments, we examined the inclusion of clathrin adaptor proteins, AP-1 and AP-2, as discrete point particles in the cytoplasmic and cytosolic interface regions. However, the approach of modelling clathrin adaptor binding as bimolecular reactions had two major limitations. First, binding of an adaptor protein to CEACAM1 does not directly transport the molecule to its destination. Instead, transport requires a host of additional proteins to recruit clathrin and promote the formation of vesicles containing multiple copies of CEACAM1 [33]. Second, following assumptions for Smoluchowski reactions, reactions in Cell4D occur at rates that are diffusion-limited. Given that vesicles form over a timescale of the order of seconds, their formation violates this assumption. Although Cell4D could simulate sufficient CCV transport events to reach system equilibrium, an implementation of these events would require simulations lengths of hundreds of millions of timesteps, requiring extended computational run-times. As an example, the CEACAM1 lipid raft model described above were performed for 200,000 timesteps, taking approximately 5 hours on a single Intel 80- thread CPU running at 2.4GHz.

Computational models can be a powerful tool for understanding biological systems. Ideally, models maintain a balance where details that have a low impact on system behaviour are ignored or abstracted away, while more critical aspects are preserved, producing models robust to input parameters and capable of emergent behaviors that provide novel insights and testable hypotheses. Such tools, we argue, are best constructed and interpreted by cell biologists who are the domain experts. For this we built Cell4D to be a user-friendly, feature-rich, and flexible framework for users to develop complex pathway models and generate large amounts of simulation data. Further, we developed a graphic interface to allow visualization of Cell4D simulations in real time. Such visualizations enhance a user’s understanding of the system being modeled which is especially useful when observing the impact of changing parameters. For example, visualization of our CEACAM1 system allows users to see how increasing concentrations of calmodulin result in increased surface accumulation of CEACAM1. Examples of simulation visualizations are provided as movies on the project GitHub site: https://github.com/ParkinsonLab/cell4d.

## ACKNOWLEDGEMENTS

Thanks to D. Han and K. Keerthisingham for additional coding support and N. Liu for software testing. We are grateful to A. Funahashi for advice on XML schema.

## Supplemental Files

Supplemental File 1: Cell4D XML Schema

Supplemental Text: Detailed Methods and Supplemental Figure Legends

