## Supplementary figures and images for "Cell4D: A general purpose spatial stochastic simulator for cellular pathways"

### Supplemental Figure 1

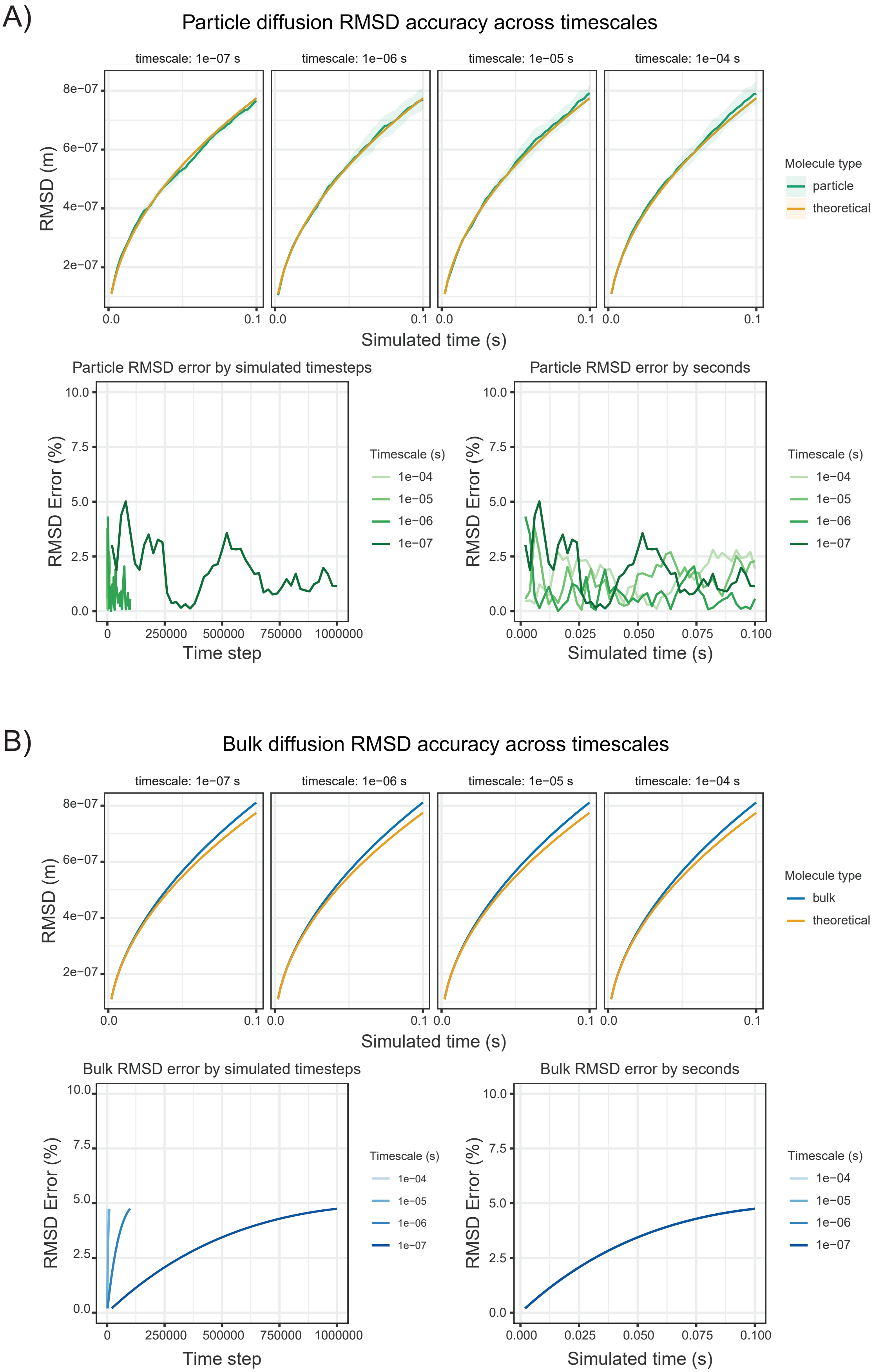

### Supplemental Figure 2

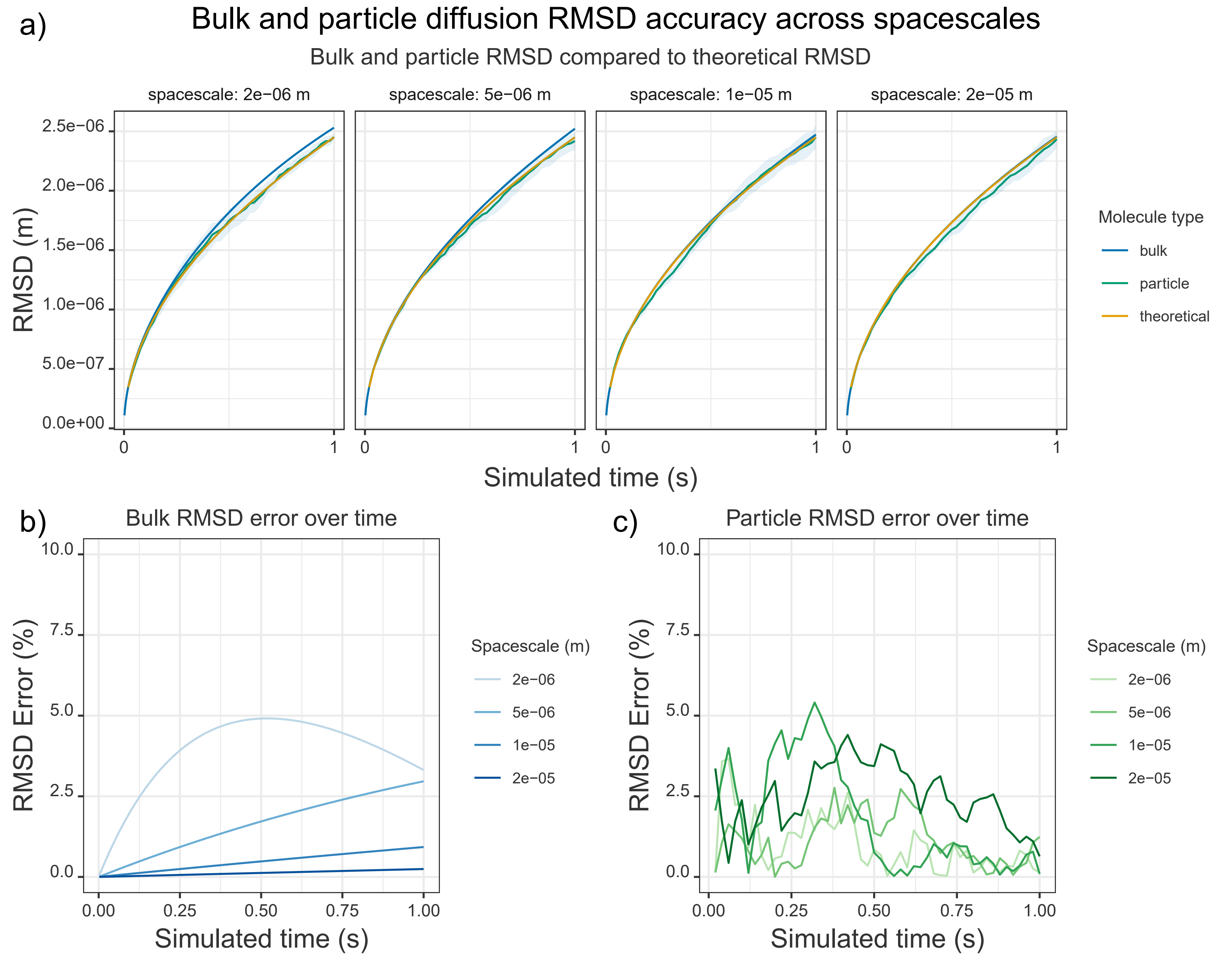

### Supplemental Figure 3

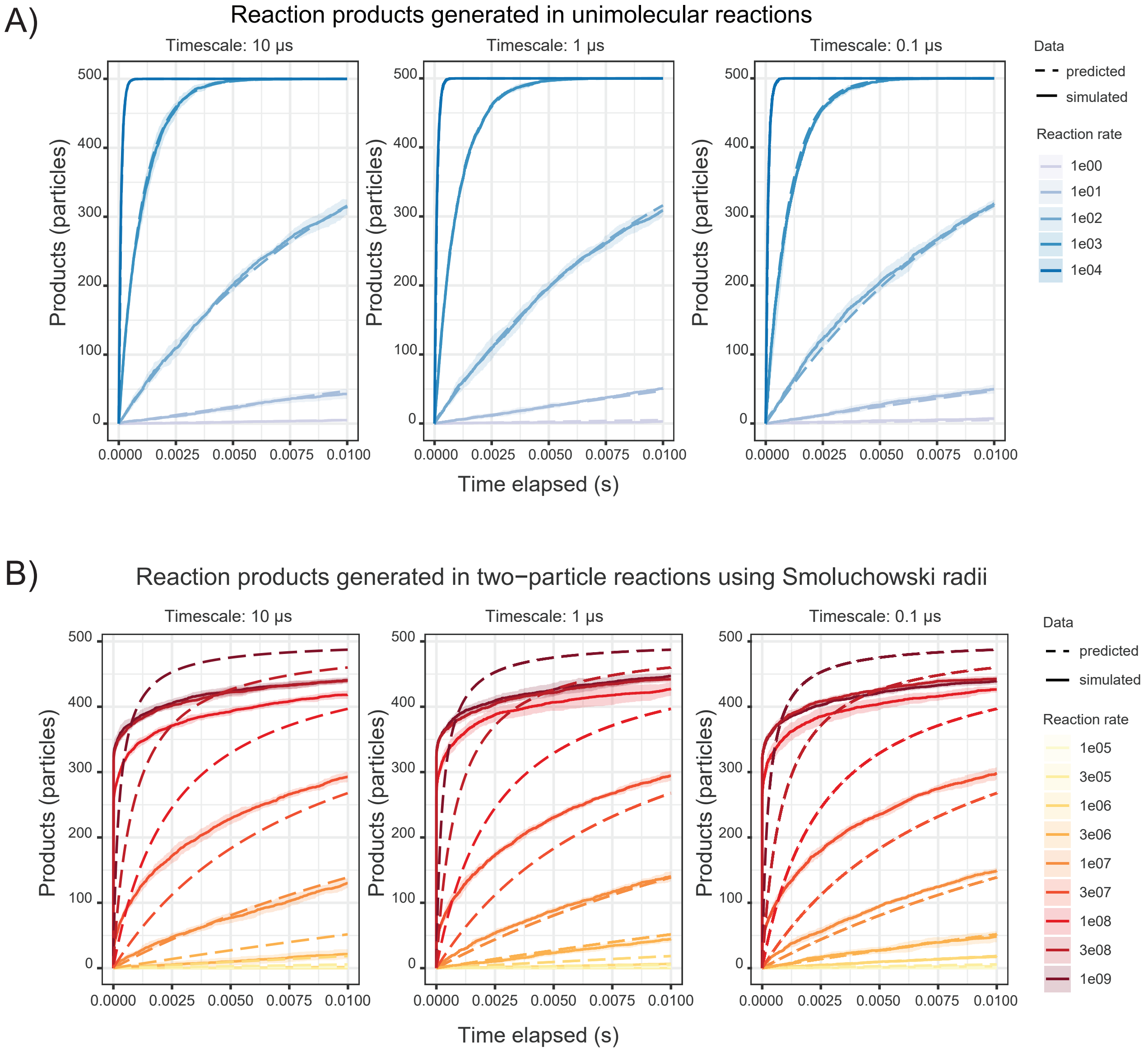

### Supplemental Figure 4

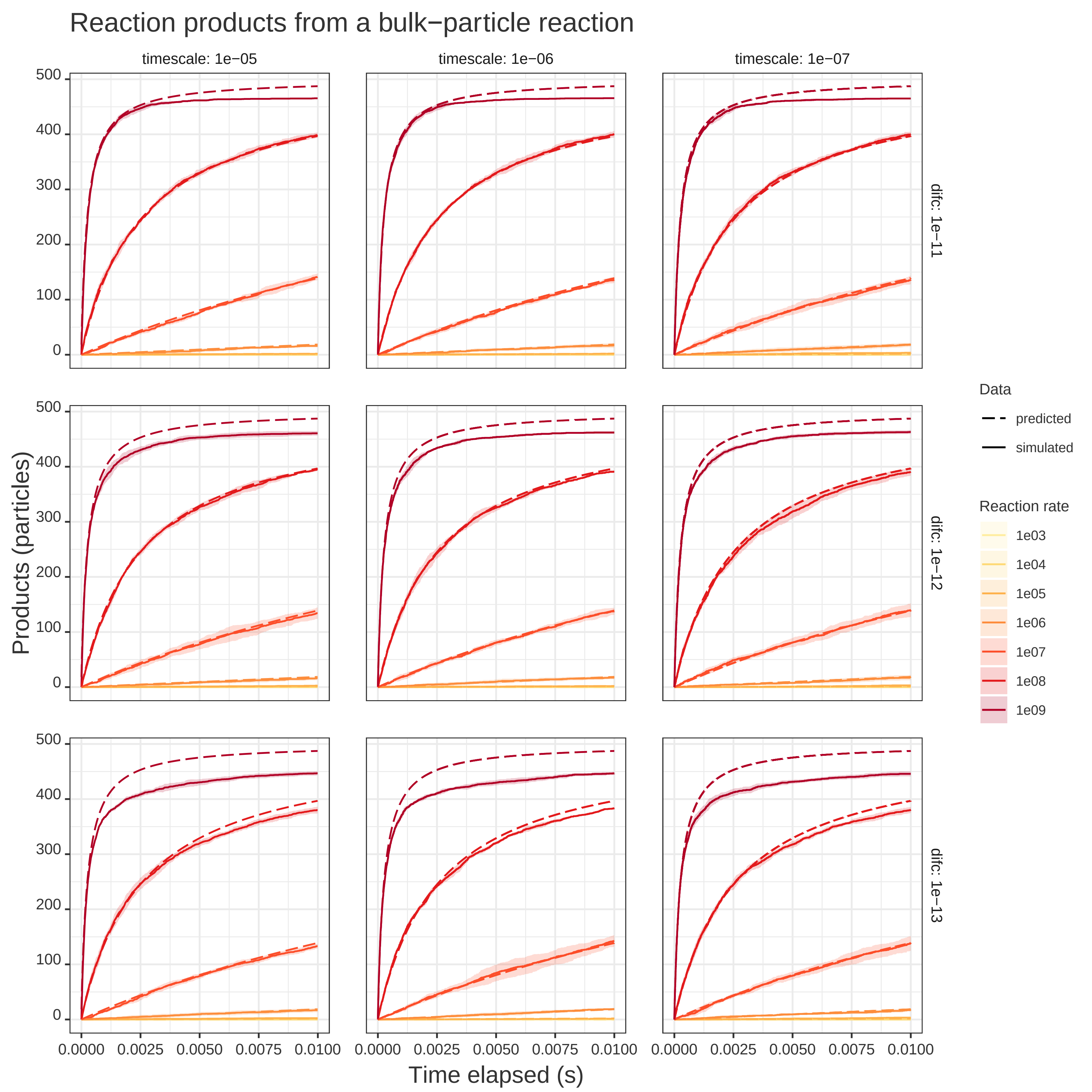

### Supplemental Figure 5

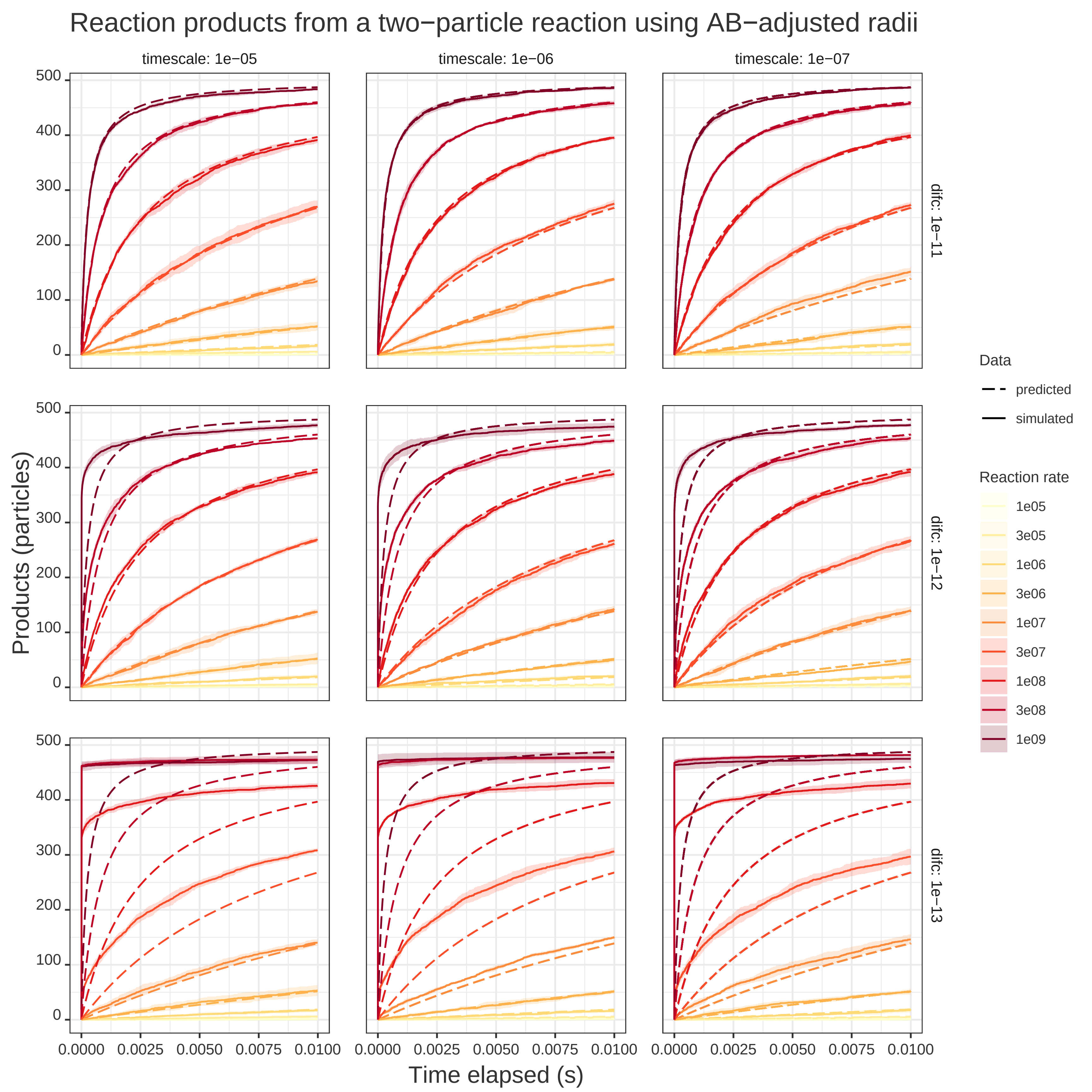

### Supplemental Figure 6

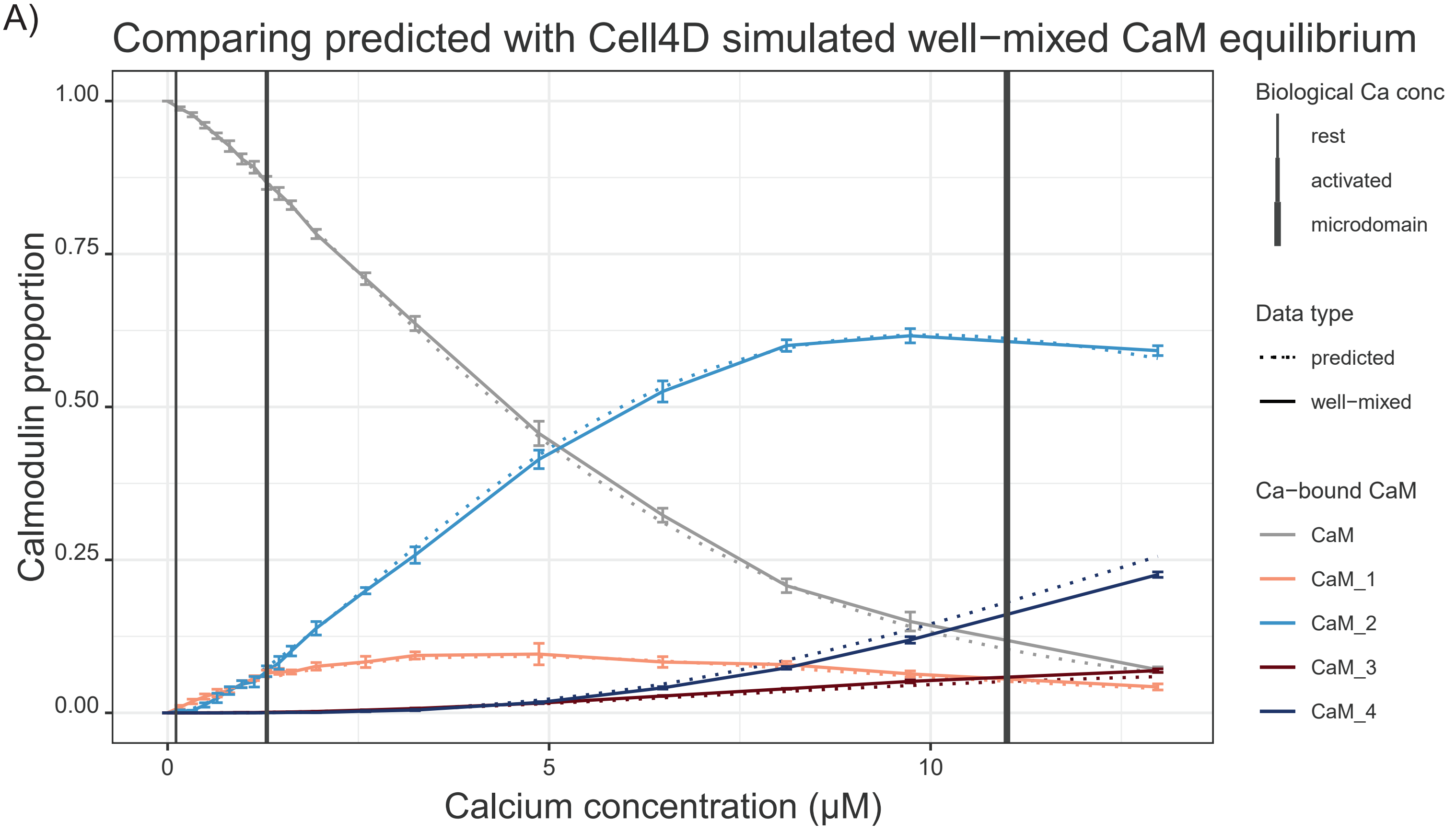

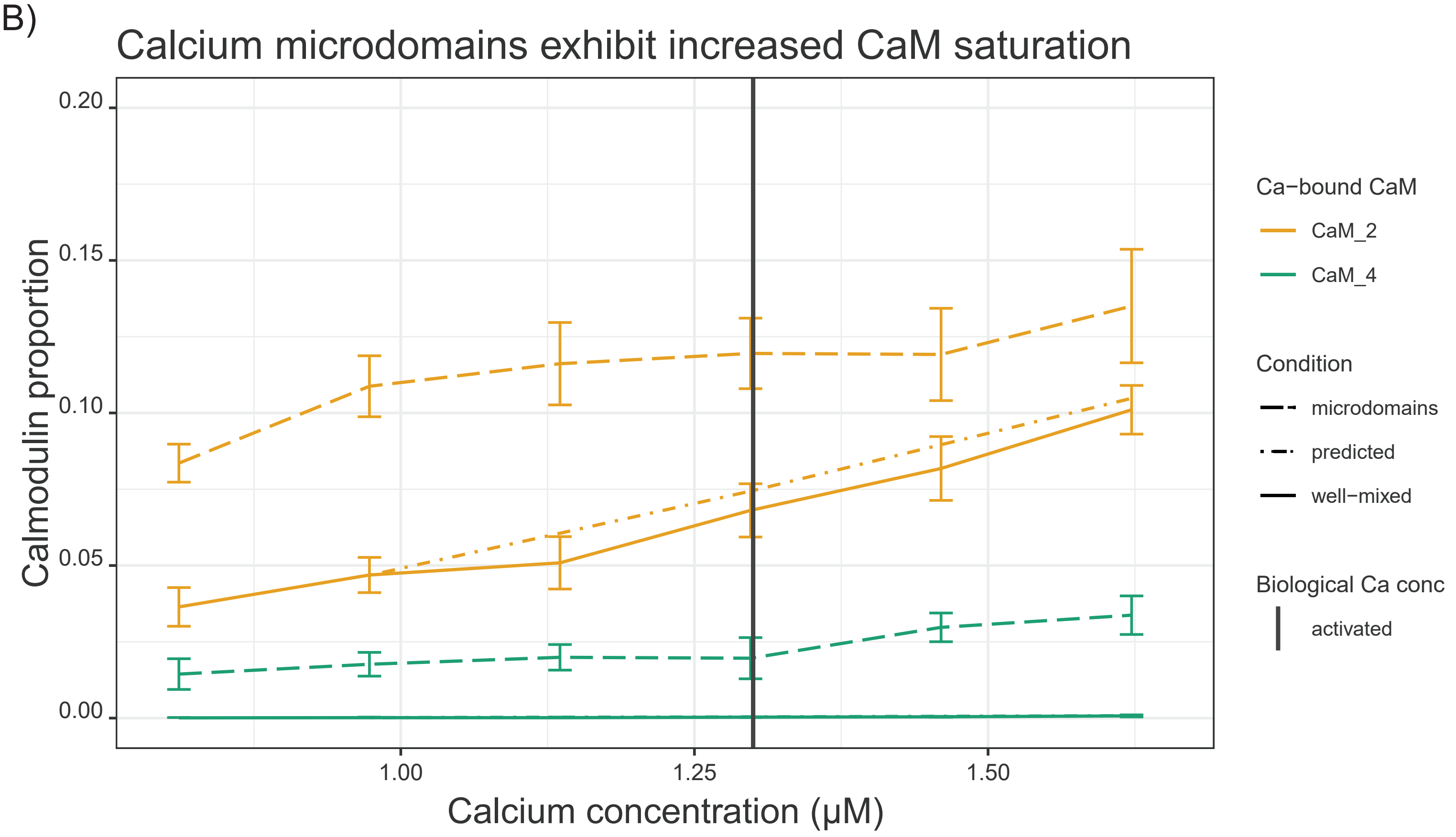

### Supplemental Figure 7

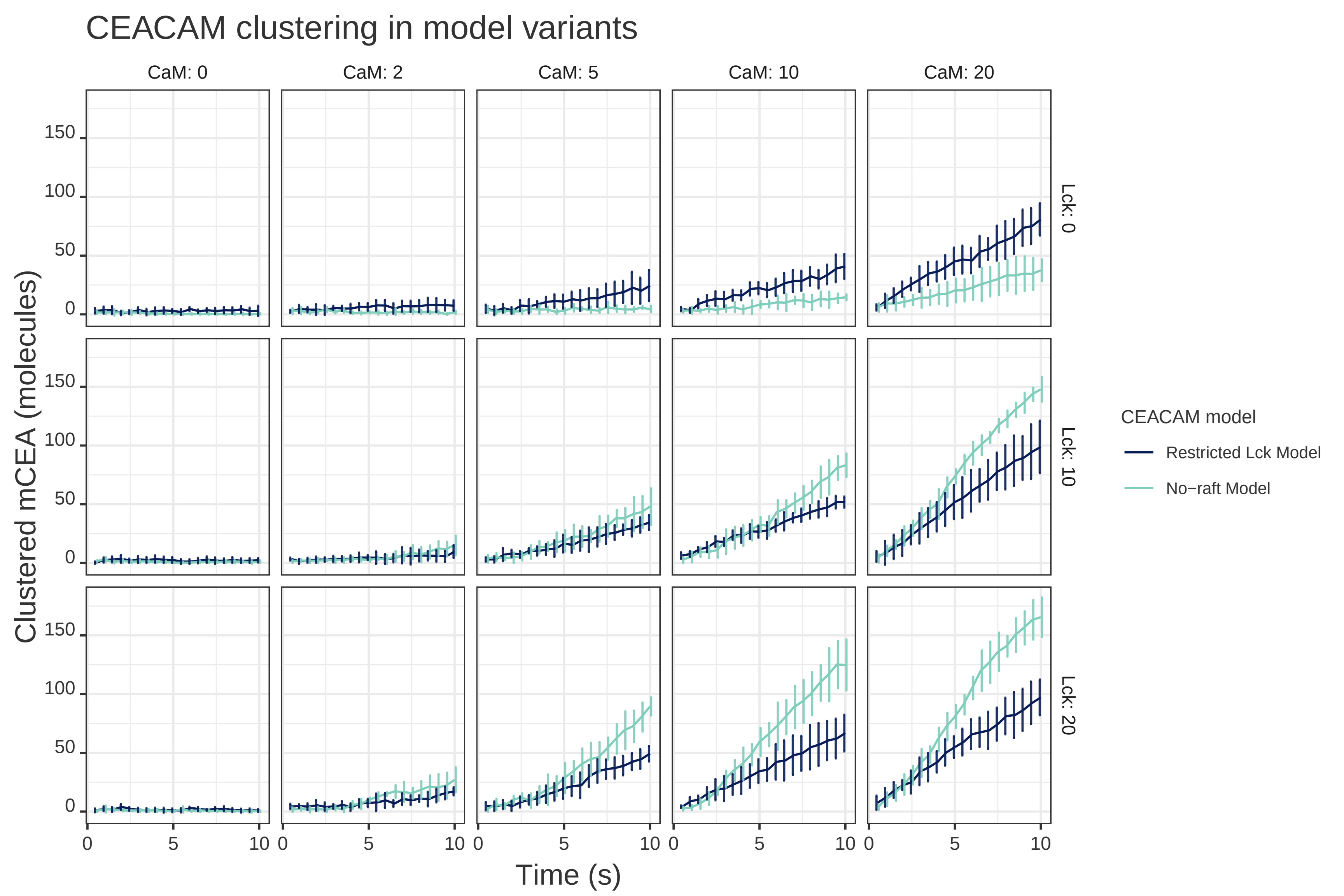
