## Supplemental Text for "Cell4D: A general purpose spatial stochastic simulator for cellular pathways"

**Additional details of System and Methods**

***Data model***

Cell4D models are stored as XML documents and are compliant with systems biology markup language (SBML version 2) (Finney and Hucka, 2003) (See schema: Supplementary File 1: Cell4D.xsd). For additional functions, not supported in SBML, an extended schema was designed that embeds Cell4D-specific XML tags, an approach inspired by CellDesigner (Matsuoka, et al., 2014). These tags provide support for lattice-based compartments, binding sites, event triggers, rules for formation of molecular complexes, states and their transitions, molecular transport etc. and contain the following mandatory layers:

Environmental Variables - Encodes general system information such as the dimensions of simulation space, the time scale and space scale of simulation, and the number of time-cycles

Species Types - Defines the list of “base molecules” that can be combined to form larger molecular complexes.

Compartments – Specifies spatial coordinates belonging to compartments that define regions where molecular species are or are not allowed to enter by diffusion.

Species - Defines specific molecules (and complexes) inclusive of their relevant states that will be represented in the model, including their number, and starting positions.

Reactions - Defines all possible reactions in the model, with layers for the list of reactants, products, and reaction kinetics. These reactions can be limited to certain spatial compartments.

Events (optional layer) - Defines triggers (time-based or condition-based) that result in the addition or removal of molecules during the simulation or the transport of molecules from one compartment to another.

***Data Flow***

Cell4D is written in C++. Model files are loaded into the simulation environment and a random number seed is either generated or provided by the user (allows recapitulation of a simulation). The simulation then iterates over a user-defined number of timesteps. Within each timestep, molecular diffusion events are first simulated followed by reaction events (**Figure 1**). For small molecules represented by local concentrations within a lattice cell (c-voxel), the diffusion into neighboring c-voxels are calculated deterministically based on follows Fick’s law (Fick, 1855). For molecules and complexes represented as point particles, displacement is determined either stochastically (i.e., Brownian motion) or deterministically (i.e. active transport).

Reaction events begin with unimolecular reactions, in which the simulator engine first ensures the presence of valid reactants (defined through a reaction schema capturing binding or modification states), employing a probabilistic model based on the defined reaction rate. Completed reactions alter the states of reactants as defined in the reaction scheme. Products generated in a reaction are placed at the same position as the original reactant. For multi-product reactions, the products are placed randomly within a small radius of the original reactant. Next bimolecular reactions are assessed by computing the distance between reaction pairs. Products of bimolecular reactions are placed within a random radius of the midpoint of the reactive species. All reaction products are prohibited from further reactions in the same timestep, eliminating the immediate reversal of reactions. An optional active transport component can also be defined for reaction products, resulting in directed movement towards a defined destination.

Additional features for running Cell4D in a high-performance computing (HPC) environment include a “no graphics” mode and, the inclusion of checkpoints to safeguard against unexpected termination agents and facilitating simulations with extended run-times. Two types of simulation output files are generated over the course of the simulation. The first type are tab-separated text files that record the counts of each type of molecular species in each compartment at every timestep. The second file type uses JavaScript Object Notation (JSON) format to record the initial random number seed used to initialize the simulation, together with position and state data of all molecules recorded at every 10,000th timestep. In addition to reporting system dynamics, this latter file serves as the input checkpoint file for restarting simulations.

***User Interface***

The graphical simulation environment includes a real-time visualization of the running simulation as well as side-panel controls enabling the user to manipulate the speed of the simulation (timesteps/sec), rotate and zoom the simulation, and select subsets of molecules and compartments to highlight in the display. Intuitively, the same selections are used to graph molecule frequencies over time.

***Model construction***

To help construct valid XML model files, we provide a web-based application – Cell4D Model Editor - accessible at http://compsysbio.org:3001. Written in React JS, tabbed sections provide prompts for the necessary and optional inputs describing each layer of the data model. The input is validated against the Cell4D schema, ensuring that data dependencies are met before the model is saved. Decoupling the model builder from the simulator allows for customization without necessarily involving the simulator code e.g., adding comment text or other metadata to the model file or changing the look and feel of the model builder.

**Additional details of Algorithms**

***Point particle movement***

The random diffusion of any particle for each timestep is determined by sampling from a normal distribution centered at 0 and a standard deviation of:

$\sqrt{2D_{B}\Delta t}$	[1]

for each cartesian coordinate, where D_B_ represents the diffusion coefficient of the particle and ∆t is the length of the timestep. For 3D space, this results in an overall root mean squared displacement (RMSD) of:

$s=\sqrt{6D_{B}\Delta t}$	[2]

If the particle encounters an impermeable boundary (e.g., compartment boundary), they are deflected in a random direction. For partially permeable compartments, the molecule will either transmit through or be deflected from the boundary based on a transmission probability defined for that compartment.

***Bulk molecule movement***

Bulk molecules in Cell4D are modelled as concentrations within a c-voxel. Molecular diffusion for these species follows Fick’s laws of diffusion (Fick, 1855) and is simulated using the forward Euler method (Pletcher, 1997). The concentration of molecules in each c-voxel at time t^n^ is used iteratively to model the changes caused by molecule flux resulting in a subsequent model state at time t^n+1^ (**Figure 1**). Diffusion between discretized spaces under Fick’s law requires that there exists an interface of non-zero area between the two spatial volumes. Diffusion between each face-sharing neighbor is calculated for all neighbors of all c-voxels at every timestep to determine diffusive flux of molecules over time.

***Fick’s laws of diffusion***

Derived by Adolph Fick in 1855, Fick’s first and second laws of describe how mass is transported through diffusive means and is analogous to other relationships such as Fourier’s law, describing heat, and Ohm’s law, describing the movement of electric charge (2006; Fick, 1855).

$J=-D\frac{dc}{dx}$	[3]

Fick’s first law describes diffusive flux of a substance through an area over time, J, with the units of substance (moles or particles) divided by meters squared time. D represents the diffusion constant of the substance in question, in units of meters squared over time, c is the concentration, and x is distance travelled. An interpretation of this equation is that molecules diffuse from an area of high concentration to low concentration, and this rate is proportional to the difference in concentration between the two areas. Fick’s second law describes the change in concentration over time caused by diffusion.

$\frac{dc}{dt}=D\frac{d^{2}c}{dx^{2}}$	[4]

These two equations form the basis for how diffusion is modelled within Cell4D.

***Molecular Reactions***

Cell4D uses a rule-based simulation paradigm which avoids the need to explicitly define every potential reaction in a system. Instead, reactions are implicitly defined based on presence of substrates. For example, a reaction involving a dimer composed of two molecules of species, ***A*** (***AA***) with a single molecule of species, ***B***, will use the same reaction as a monomer of ***A***. Thus, if the monomeric equation is defined by $\boldsymbol{A}+\boldsymbol{B}\to\boldsymbol{C}$, the dimeric equation is implicitly defined as $\boldsymbol{A}\boldsymbol{A}+\boldsymbol{B}\to\boldsymbol{AC}$. In cases where such implicit rules are not desired, the user can define *states* for each species to restrict the application of implicit rules.

For point particles, Cell4D allows the definition of unimolecular and bimolecular reactions, the latter involving both other point particles and bulk molecules. For spontaneous unimolecular reactions involving point particles, the probability that a specific molecule, ***A***, undergoes a reaction with reaction rate, k, in a timestep of length Δt, is given by:

$P\left( \boldsymbol{A}reacts \right)=1-e^{-k \Delta t}$	[5]

For bimolecular reactions involving point particles, Cell4D uses the Andrews-Bray-adjusted Smoluchowski method. In brief, each reactant has a defined reaction radius which determines if a reaction can occur. The probability that the reaction occurs is defined by equation [1] with the reaction rate, k, defined as:

$k=4\pi D_{M}\sigma$	[6]

Where D_M_ is the mutual diffusion rate constant constant (m^2^/s), defined as the sum of the diffusion rates of the two reactants, and σ is the distance between the two reactants. If the reaction occurs, products are placed either at the midpoint of the reactants if not reversible, or placed outside the reaction radius (defined as 1.2 x the radium of the forward reaction radius) to avoid spontaneous reaction reversibility.

If a molecule is capable of performing more than one possible reaction during a given timestep, the system first calculates the probability of the molecule undergoing *no* reaction. If a reaction does occur, it is selected from a discrete distribution based on the rate constant of each possible unimolecular or bimolecular reaction. For bimolecular reactions, if there is more than one potential reaction partner (i.e. within the defined reaction radius), one is selected at random. Since this method is sensitive to errors where diffusion distance of molecules exceeds or is close to the reaction radius, Cell4D applies the Andrews-Bray-adjustment to artificially increase reaction radii of molecules (Andrews and Bray, 2004).

For bulk molecules, they can either undergo enzymatic reactions or participate in reactions that modulate the state of point particles (e.g. phosphorylation status). Enzymatic reactions are defined by the reversible Michaelis Menten equation:

$v= \frac{\frac{V_{\max}^{f}}{K_{m}^{s}}\left[ S \right]- \frac{V_{\max}^{r}}{K_{m}^{p}}\left[ P \right]}{1+ \frac{\left[ S \right]}{K_{m}^{s}}+ \frac{\left[ P \right]}{K_{m}^{p}}}$	[7]

where v^f^_max_ and v^r^_max_ are the maximum rates of reaction for the forward and reverse reactions respectively, and K^s^_m_ and K^p^_m_ are the Michaelis constants for the subtrate and product respectively. For state changes, Cell4D uses equations [5] and [6] with k multiplied by the number of bulk molecules in the current voxel and the time step, Δt. In cases where a particle could react with more than one type of bulk molecule within its c-voxel, the reaction is determined by the relative probability of each type of reaction.

***Integrated rate law for second-order reactions***

Since second-order reactions require collisions between molecules, this adds an additional variable, diffusion rate, to the determination of the bulk reaction rate.

$rate=\frac{d[AB]}{dt}=-\frac{d\left[ A \right]}{dt}=-\frac{d[B]}{dt}= -k \times\left[ A \right]\times\left[ B \right]$

$\left[ A \right]=\left[ B \right]={[A]}_{0}-[AB]$

$\begin{aligned} rate=-\frac{d\left[ A \right]}{\mathrm{dt}}=-k \times\left[ A \right]^{2}\# \end{aligned}$

Where [A] and [B] represent the concentration of each reactant molecule, and [AB] is the concentration of the bimolecular reaction product. The integrated rate law for a second-order reaction is more complicated, but a special case can be applied for when both reactants have the same concentration at Δt = 0 and react with the same stoichiometry, which is the case used in the validation simulations.

$\int_{{[A]}_{0}}^{{[A]}_{\Delta t}} \frac{d[A]}{\left[ A \right]^{2}}=-\int_{0}^{\Delta t} k dt$

$\frac{1}{{[A]}_{\Delta t}}=kt+\frac{1}{\left[ A \right]_{0}}$

The integrated rate law derived above is used to calculate the formation of products over time then compared to the results from corresponding Cell4D simulations using the same reaction parameters.

*Smoluchowski model for diffusion-limited reactions*

In 1917, Marian Smoluchowski presented a mathematical description of reaction rates on the scale of individual molecules, postulating that reactions occur when two reactants are sufficiently close to each other, introducing the influence of diffusion in reactions. In the absence of diffusion, the laws of mass action kinetics predict that reactions with a sufficiently large K would have reactants instantly react once they are in proximity, and all possible reactants will instantaneously be converted to products. Instead, in his model Smoluchowski proposed a diffusion-controlled reaction rate, K_S_ which can replace the mass action kinetic rate K when K is sufficiently large.

$\begin{aligned} K_{s}=4\pi D_{M}\sigma\# \end{aligned}$

The reaction rate of these kinetically fast reactions is defined by the Smoluchowski reaction rate, K_S_, which depends on D_M_, the diffusion constant (m^2^/s) of the reactants, and σ, the binding radius where a reaction would occur if the reacting molecules are within that distance. The Collins-Kimball relation shows that the effective rate of a reaction is essentially the same as the diffusion-limited Smoluchowski rate, K_S_, when K ≫ K_s_ (Collins and Kimball, 1949).

$\begin{aligned} \frac{1}{K_{\mathrm{eff}}}=\frac{1}{K}+\frac{1}{K_{S}}\# \end{aligned}$

Thus, under the Smoluchowski scheme, all bimolecular reactions are assumed to be diffusion-controlled, and their rates will be calculated using the above K_s_ equation. With the exception of enzymatic reactions calculated using methods based on reversible Michaelis-Menten kinetics, all bimolecular reactions within Cell4D are modelled based on Smoluchowski’s diffusion-controlled reaction equation.

***Implementation of Andrews-Bray correction of Smoluchowski reaction radii for Cell4D bimolecular reactions***

There are two methods that Cell4D uses to convert input reaction rates into binding radii that the simulator can use. The first method is using the Smoluchowski equation (see previous section), reversing the equation to calculate the corresponding reaction radii using the input rate constant combined with the corresponding diffusion rates of the reactants. The second method was developed by Andrews and Bray for the simulation program Smoldyn, which uses a modified Smoluchowski approach (AB method; (Andrews and Bray, 2004)). A smoothed version of the look-up table of values provided with Smoldyn is implemented by Cell4D. The input value of the lookup table is the “reduced step length,” *s’* and the output is the “reduced reaction rate,” *k’*, which are the respective average step length of a particle within a timestep and the experimental reaction rate that are normalized by the binding radius of the reaction to create unitless values.

$s^{'}=\frac{s}{\sigma_{b}}$

$k^{'}=\frac{k \Delta t}{{\sigma_{b}}^{3}}$

Using the look-up table, the corresponding (reduced) reaction rate can be found if the reduced step length of the reaction is known, which requires knowledge of both the mutual step length of the reacts as well as the theoretical binding radius.

This solves the forward problem where a rate constant can be found from known reaction radii and diffusion constants, but here the reverse problem needs to be solved, because the binding radius must first be known to utilize the look-up table. To solve this reverse problem, an iterative procedure is implemented (after (Andrews and Bray, 2004)) to estimate the correct binding radius based on the original Smoluchowski equation. After initially estimating the binding radius, the reduced step length is calculated and used to reference the look-up table for the reduced reaction rate. This rate is then converted back to obtain an estimated rate constant. This rate constant is then compared to the original input rate constant, and the error is used for the next estimate of the binding radius. This process is repeated until the estimated radius produces an estimated rate that is within 0.001% of the input rate constant.

Since the look-up table has a finite number of possible values, where reduced step lengths ranging from e^-3^ to e^3^ are paired with their corresponding reduced reaction rate constants, we fit the 31 data points of the table with a 5-parameter log-logistic curve in R using the “drc” package (R Development Core Team, 2010; Ritz, et al., 2016). This equation is used within the Cell4D to iteratively arrive at the estimated radius of reactions using the method outlined above. Since the solution from this method approaches the base Smoluchowski method for reactions with a low reduced step length, we developed a program that automatically uses the unmodified Smoluchowski equation to calculate the radius if the reduced step length is lower than the lowest value pair provided in the table. This avoids unnecessary extrapolation from the table creating inaccurate reaction radius predictions. The function (*calculate_AB_radius*) is available in the simulation code on the project GitHub site: https://github.com/ParkinsonLab/cell4d.

In the AB-adjusted reduced rate determination, Cell4D implements the function:

$$\begin{aligned} {rate}_{r}=c+ \frac{d-c}{\left( 1+\exp\left( b\times\left( \log\left( s_{r} \right)-e \right) \right) \right)^{f}}\# \end{aligned}$$

Where:

$$\begin{aligned} {rate}_{r}=\frac{k\times\Delta t}{{\sigma_{b}}^{3}}\# \end{aligned}$$

$$\begin{aligned} s_{r}=\frac{\sqrt{2\times\Delta t\times D_{M}}}{\sigma_{b}}\# \end{aligned}$$

To improve accuracy, two versions of the function are used, with different parameters for the five variables, depending on the step lengths of the reactants.

For reduced step length (s_r­_) < 1:

$b=-2.35948390$

$c = -0.00125267$

$d=4.78882764$

$e=0.00154475$

$f=0.80220128$

For reduced step length (s_r­_) > 1:

$b=-2.89171179$

$c=-0.00648886$

$d=4.19026581$

$e=0.00279511$

$f=0.62295397$

This AB adjustment is not applied to bulk-particle reaction interactions. The adjusted method accounts for the limited spatial resolution associated with finite time-steps, particularly in cases where straight line diffusion of particles might result in the transit of particles through the reaction radius of a potential reactant, potentially resulting in missed reactions. The bulk-particle interactions are partially deterministic, as the reaction probability is proportional to the concentration of the well-mixed bulk reactants within the c-voxel and adjusted by the timescale of the simulation. This means that the predicted probability of these reaction is unaffected by the limitations of simulating the reaction dynamics of two discrete particles and will predict the correct rates independent of step length and timescale.

***Molecular Complexes***

In Cell4D, each point particle in the simulation can represent more than one discrete molecule as a complex. Reactions that generate complexes such as dimers will combine the two reacting molecules into a single point particle. This approach reduces computational overhead by eliminating superfluous molecule generation/destruction, allows complexes to have derived properties that are based on their components, and leads to the possibility of flexible oligomeric or polymeric complexes that do not artificially increase the number of species variants within the simulation.

***Modeled Systems***

Calmodulin Activation

Calmodulin (CaM) is a calcium-binding protein (CBP) that consists of two Ca^2+^ binding lobes with two EF-hand motifs each, which can bind Ca^2+^ ions to induce conformational changes that promote cooperative binding properties. Each lobe possesses different binding properties. The N-terminal domain acts as a fast binding site, with a log association rate constant (log_10_k_on_) of approximately 9 M^-1^ s^-1^, while the C-terminal site binds an order of magnitude slower, log_10_k_on_ ~8 M^-1^ s^-1^. However, the fast N-terminal domain exhibits a lower affinity due to its higher log dissociation constant compared to the C-terminal domain (5.2 s^-1^ compared to 3.4 s^-1^). Thus, the N-terminal Ca^2+^ sites allow calmodulin to rapidly bind calcium ions, acting as a calcium sink, while the higher dissociation rate enables Ca^2+^ binding with the slower, yet higher affinity C-terminal sites.

In addition, differential binding at the two types of lobes, CaM also exhibits co-operative binding with calcium. In this system, each lobe acts independently. For each lobe, each of the binding domains can be represented by two forms: the unbound tight (T) form, and the Ca^2+^-bound relaxed (R) form. Ca^2+^ binding for each lobe is subsequently modeled as a two step system (Faas, et al., 2011).

$N_{T}N_{T}+{Ca}^{2+}\begin{matrix} 2\times k_{on\left( T \right),N} \\ \leftrightarrow\\ k_{off\left( T \right),N} \end{matrix}CaN_{T}N_{R}+{Ca}^{2+}\begin{matrix} k_{on\left( R \right),N} \\ \leftrightarrow\\ 2\times k_{off\left( R \right),N} \end{matrix}CaN_{R}{CaN}_{R}$

$C_{T}C_{T}+{Ca}^{2+}\begin{matrix} 2\times k_{on\left( T \right),C} \\ \leftrightarrow\\ k_{off\left( T \right),C} \end{matrix}CaC_{T}C_{R}+{Ca}^{2+}\begin{matrix} k_{on\left( R \right),C} \\ \leftrightarrow\\ 2\times k_{off\left( R \right),C} \end{matrix}CaC_{R}CaC_{R}$

Here, both sites within a specific terminus are considered equivalent, and the first step of the binding will have twice the rate of k_on(T)_ since both binding sites are valid. The reverse reaction will have the normal k_off(T)_ value since only one Ca2+ is bound to the CaM. For step 2, calcium binding to the remaining site will have a rate of k_on(R)_, while the reverse reaction, the dissociation of either of the calcium ions from the relaxed state, will be twice the rate of k_off(R)_. Using this scheme, only 8 parameters are necessary to model calcium-CaM interactions. Experimental rate constants of these reactions have been determined previously through Ca2+ uncaging measurements with wild-type calmodulin as well as mutants with a single non-functional binding

Terminus (Faas, et al., 2011).

In terms of cooperativity, the two termini exhibit the same effect of increased Ca^2+^ upon binding of one of the sites, but the mechanisms employed are different. For the N-terminus, the association and dissociation rates exhibit an order of magnitude increase and decrease respectively during a T to R shift (log_10kon_ 8.9 to 10.5 M^-1^s^-1^, log_10koff_ 5.2 to 4.3 s^-1^). On the other hand, the C-lobe association rate does not change dramatically (log_10kon_ 7.9 to 7.4 M^-1^s^-1^), but the dissociation rate decreases by over 100-fold (log_10koff_ 3.4 to 1.2 s^-1^). This means that the Ca^2+^ affinity of fully loaded CaM is higher than the affinity of unbound CaM, contributing to the observed switch-like behaviour.

*Deterministic Euler forward CaM equilibrium equations*

K_on(1),C_ = 8.4 × 10^7^

K_on(2),C_ = 2.5 × 10^7^

K_off(1),C_ = 2.6 × 10^3^

K_off(2),C_ = 6.5 × 10^0^

K_on(1),N_ = 7.7 × 10^8^

K_on(1),N_ = 3.2 × 10^10^

K_on(1),N_ = 1.6 × 10^5^

K_on(1),N_ = 2.2 × 10^4^

$$\frac{\Delta{[CaM}_{N1}]}{\Delta t}= {(2 \times k}_{on\left( 1 \right),N}\times\left[ CaM \right]\times\left[ {Ca}^{2+} \right]+k_{off\left( 1 \right),N} \times\left[ {CaM}_{N1C1} \right]+ 2\times k_{off\left( 2 \right),N} \times[{CaM}_{N2}])-{(2 \times k}_{on\left( 1 \right),C}\times\left[ {CaM}_{N1} \right]\times[{Ca}^{2+}]+k_{on\left( 2 \right),N} \times[{CaM}_{N1}] \times[{Ca}^{2+}]+ k_{off\left( 1 \right),N} \times[{CaM}_{N1}])$$

$$\frac{\Delta{[CaM}_{C1}]}{\Delta t}= {(2 \times k}_{on\left( 1 \right),C}\times\left[ CaM \right]\times\left[ {Ca}^{2+} \right]+k_{off\left( 1 \right),C} \times\left[ {CaM}_{N1C1} \right]+ 2\times k_{off\left( 2 \right),C} \times[{CaM}_{C2}])-{(2 \times k}_{on\left( 1 \right),N}\times\left[ {CaM}_{C1} \right]\times[{Ca}^{2+}]+k_{on\left( 2 \right),C} \times[{CaM}_{C1}] \times[{Ca}^{2+}]+ k_{off\left( 1 \right),C} \times[{CaM}_{C1}])$$

$$\frac{\Delta{[CaM}_{N2}]}{\Delta t}= {(k}_{on\left( 2 \right),N}\times\left[ {CaM}_{N1} \right]\times\left[ {Ca}^{2+} \right]+k_{off\left( 1 \right),N} \times\left[ {CaM}_{N2C1} \right])-{(2 \times k}_{on\left( 1 \right),C}\times\left[ {CaM}_{N2} \right]\times[{Ca}^{2+}]+k_{on\left( 1 \right),C} \times[{CaM}_{N2}] \times[{Ca}^{2+}])$$

$$\frac{\Delta{[CaM}_{C2}]}{\Delta t}= {(k}_{on\left( 2 \right),C}\times\left[ {CaM}_{C1} \right]\times\left[ {Ca}^{2+} \right]+k_{off\left( 1 \right),C} \times\left[ {CaM}_{N1C2} \right])-{(2 \times k}_{on\left( 1 \right),N}\times\left[ {CaM}_{C2} \right]\times[{Ca}^{2+}]+k_{on\left( 1 \right),N} \times[{CaM}_{C2}] \times[{Ca}^{2+}])$$

$$\frac{\Delta{[CaM}_{N1C1}]}{\Delta t}=(2\times k_{off\left( 2 \right),C} \times\left[ {CaM}_{N1C2} \right]+ 2\times k_{off\left( 2 \right),N} \times[{CaM}_{N2C1}] + 2 \times k_{on\left( 1 \right),N}\times\left[ {CaM}_{C1} \right]\times\left[ {Ca}^{2+} \right]+ {2 \times k}_{on\left( 1 \right),C}\times\left[ {CaM}_{N1} \right]\times\left[ {Ca}^{2+} \right])-(k_{on\left( 2 \right),N}\times\left[ {CaM}_{N1C1} \right]\times[{Ca}^{2+}]+k_{on\left( 2 \right),C}\times\left[ {CaM}_{N1C1} \right]\times\left[ {Ca}^{2+} \right]+ k_{off\left( 1 \right),N} \times\left[ {CaM}_{N1C1} \right] + k_{off\left( 1 \right),C} \times\left[ {CaM}_{N1C1} \right])$$

$$\frac{\Delta{[CaM}_{N1C2}]}{\Delta t}=(2 \times k_{on\left( 1 \right),N}\times\left[ {CaM}_{C2} \right]\times\left[ {Ca}^{2+} \right]+ k_{on\left( 2 \right),C} \times\left[ {CaM}_{N1C1} \right]\times\left[ {Ca}^{2+} \right]+ 2\times k_{off\left( 2 \right),N} \times[{CaM}_{N2C2}])-(k_{on\left( 2 \right),N}\times\left[ {CaM}_{N1C2} \right]\times[{Ca}^{2+}]+ 2 \times k_{off\left( 2 \right),C} \times\left[ {CaM}_{N1C2} \right]+ k_{off\left( 1 \right),N} \times\left[ {CaM}_{N1C2} \right])$$

$$\frac{\Delta{[CaM}_{N2C1}]}{\Delta t}=(2 \times k_{on\left( 1 \right),C}\times\left[ {CaM}_{N2} \right]\times\left[ {Ca}^{2+} \right]+ k_{on\left( 2 \right),N} \times\left[ {CaM}_{N1C1} \right]\times\left[ {Ca}^{2+} \right]+ 2\times k_{off\left( 2 \right),C} \times[{CaM}_{N2C2}])-(k_{on\left( 2 \right),C}\times\left[ {CaM}_{N2C1} \right]\times[{Ca}^{2+}]+ 2 \times k_{off\left( 2 \right),N} \times\left[ {CaM}_{N2C1} \right]+ k_{off\left( 1 \right),C} \times\left[ {CaM}_{N2C1} \right])$$

$$\frac{\Delta{[CaM}_{N2C2}]}{\Delta t}=(k_{on\left( 2 \right),C} \times\left[ {CaM}_{N2C1} \right]\times\left[ {Ca}^{2+} \right]+ k_{on\left( 2 \right),N} \times\left[ {CaM}_{N1C2} \right]\times\left[ {Ca}^{2+} \right])-(2 \times k_{off\left( 2 \right),N} \times\left[ {CaM}_{N2C2} \right]+ 2 \times k_{off\left( 2 \right),C} \times\left[ {CaM}_{N2C2} \right])$$

$$\left[ CaM \right]=\left[ CaM \right]_{0}-(\left[ {CaM}_{N1} \right]+ \left[ {CaM}_{N2} \right]+ \left[ {CaM}_{C1} \right]+ \left[ {CaM}_{C2} \right]+\left[ {CaM}_{N1C1} \right]+ \left[ {CaM}_{N1C2} \right]+ \left[ {CaM}_{N2C1} \right]+ \left[ {CaM}_{N2C2} \right])$$

$$\left[ Ca \right]=\left[ Ca \right]_{0}-\left( \left[ {CaM}_{N1} \right]+\left[ {CaM}_{C1} \right] \right)-2\times\left( \left[ {CaM}_{N2} \right]+\left[ {CaM}_{C2} \right]+\left[ {CaM}_{N1C1} \right] \right)-3\times\left( \left[ {CaM}_{N1C2} \right]+\left[ {CaM}_{N2C1} \right] \right)-4\times(\left[ {CaM}_{N2C2} \right])$$

Models focused on calmodulin activation used a simulation environment defined as a cube with side length of 8 × 10^-7^ m. Calcium ions were defined as bulk molecules and calmodulin was defined as a point particle with N-terminal (CaM_N) and C-terminal (CaM_C) calcium binding sites capable of representing the low-affinity unbound tense state (T) and high affinity relaxed (R) state within each binding lobe. Binding dynamics were modeled according to a two-step reaction scheme (Faas, et al., 2011). Diffusion rates of Ca^2+^ and calmodulin were set at 5.3 × 10^-10^ m/s^2^ and 1.0 × 10^-12^ m/s^2^ respectively (Donahue and Abercrombie, 1987; Sanabria, et al., 2008). The k_on(T)_ and k_off(R)_ rates were set at twice the experimental values due to the two possible calcium binding sites of each lobe. Other paramteres and equations used for the following models are detailed above.

*Well-mixed model:* A simulation timestep length of 2.5 µs was used and simulations performed for 40,000 timesteps (0.1 seconds). To account for initial stochasticity of binding events, only the final 200 timesteps (0.5 ms) were used for data analysis. Calcium was confined to the simulation environment which was defined as a single c-voxel.

*Microdomain model*: The model space was divided into a lattice comprising 10 x 10 x 10 c-voxels which were subdivided into five 2 x 10 x 10 cross-sectional compartments (quintiles Q1 thru Q5). Four c-voxels in Q1 were defined as sites of calcium release, with calcium ions introduced to maintain a concentration of 11 µM in each microdomain. The impact of three spatial configurations of calcium release were explored: 1) the four c-voxels were arranged as a 2x2 square in one corner of Q1; 2) the four c-voxels were arranged as a 2x2 square in the center of Q1; and 3) the four voxels were placed at four non-adjacent co-ordinates (3,3),(3,8),(8,3),and (8,8) in Q1. Simulations were performed with 2 µM calmodulin (616 molecules randomly placed within the simulation environment). Calcium ions were allowed to diffuse within the simulation environment and were removed if they reached the distal layer of Q5. Simulations were performed for 125,000 timesteps (0.5 seconds).

CEACAM1 activation

This model was performed within an environment composed of 6 × 6 × 6 c-voxels of length 0.2 μm. Simulations were performed for 200,000 timesteps of 50 μs (10 seconds total). The simulation environment was subdivided into: 1) an outer 1 x 6 x 6 layer, representing the ‘membrane’; 2) an adjacent 1 x 6 x 6 cytosolic interface region that allows calmodulin to interact with membrane CEACAM1 molecules; 3) a 2 x 6 x 6 ‘cytosol’ region adjacent to the cytosolic interface; and 4) a 2 x 6 x 6 ‘organelle’ representing a sink for CEACAM1 molecules. To reduce computational overhead, particles representing discrete molecules of calmodulin were confined to the cytosolic interface region. Particles located within the ‘membrane’ were constrained to 2-dimensional movement; movement of particles representing CEACAM1 molecules were allowed to enter/leave the ‘membrane’ through defined endocytosis and exocytosis transport events. For some simulations exploring the impact of ‘lipid-rafts’, the ‘membrane’ was subdivided into microdomains. These microdomains were arbitrarily defined with the co-ordinates: (0,1),(1,3),(1,4),(3,1),(4,1) and (3,5). Lck molecules with the capacity to phosphorylate CEACAM1 preventing its transport out of the membrane, when included in the model, were restricted to the membrane. Within the membrane, CEACAM1 dimers preferentially remain outside lipid-raft microdomains. Once activated (i.e. all four sites occupied with Ca^2+^), calmodulin can disassociate CEACAM1 dimers into monomers which preferentially associate with lipid-raft microdomains. At any time (within or outside lipid-raft microdomains), CEACAM1 monomers may undergo a spontaneous and reversible first-order reaction that immobilizes the CEACAM1 protein on the membrane compartment, mimicking extracellular trans-binding. Unbound CEACAM1 monomers are allowed to associate with a bound, immobilized, CEACAM1 monomer through a bimolecular reaction based on their mutual proximity. Together these two reactions allow the formation of monomeric CEACAM1 clusters on the cell surface. Model files are provided in the project GitHub: https://github.com/ParkinsonLab/cell4d.

Parameters used in these simulations are:

*Cell4D CEACAM1 model parameters*

| **General Parameters** |  |
| --- | --- |
| Timescale^1^ | 50 μs |
| Spacescale^1^ | 0.2 μm/c-voxel (1.73 × 10^-15^ L total) |
| C-voxel lattice^1^ | 6 × 6 × 6 |
| Simulation time^1^ | 10 s |
| Endocytosis rate^2^ | 325/s |
| Exocytosis rate^2^ | 118/s |

| **Compartments** | **Restricted Lck model** | **Unrestricted Lck model** | **No-raft model** |
| --- | --- | --- | --- |
| Lipid-ordered area^3^ | 2.4 × 10^-13^ m^2^ | 2.4 × 10^-13^ m^2^ | 0 |
| Lipid-disordered area^3^ | 1.2 × 10^-12^ m^2^ | 1.2 × 10^-12^ m^2^ | 1.44 × 10^-12^ m^2^ |
| CEA compart volume^1^ | 5.76 × 10^-16^ L | 5.76 × 10^-16^ L | 5.76 × 10^-16^ L |
| Lck localization regions | Lipid-ordered | Lipid-disordered  Lipid-ordered | Lipid-ordered |

| **Molecules** | **CEACAM1** | **Lck** | **Calmodulin** |
| --- | --- | --- | --- |
| Count^1^ | 510 | 0 – 20 | 0 – 20 |
| Diffusion constant (m^2^/s) ^4^ | 3×10^-13^ | 1.3×10^-12^ | 1×10^-12^ |
| Compartments | Lipid-disordered  Lipid-ordered  CEA compartment | (Lipid-disordered)  Lipid-ordered | Cytosol |

| **Reactions** | **Reactants** | **Products** | **Rate/radius** |
| --- | --- | --- | --- |
| CEA2_CaM^5^ | CEACAM1 dimer  Calmodulin | CEACAM1  CEACAM1  Calmodulin | 1 × 10^7^ M^-1^s^-1^ |
| CEA_CEA^5^ | CEACAM1  CEACAM1 | CEACAM1 dimer | 2 × 10^4^ M^-1^s^-1^ |
| CEA_trans^1^ | CEACAM1 | CEACAM1 | 2 × 10^1^ s^-1^ |
| CEA_untrans^1^ | CEACAM1 | CEACAM1 | 2 × 10^2^ s^-1^ |
| CEA_cluster^1^ | CEACAM1  CEACAM1 | CEACAM1  CEACAM1 | 2 × 10^-8^ m |
| CEA_phos^1^ | CEACAM1  Lck | CEACAM1  Lck | 2 × 10^5^ M^-1^s^-1^ |

^1^ – Model-specific parameters set arbitrarily, insensitive parameters that produces similar model outcomes across a range of possible input values

^2^ – Rates were arbitrarily set at a rate exceeding physiologically relevant vesicle-formation speeds for computational efficiency

^3^ – Lipid-ordered region surface area arbitrarily set to be 1/6^th^ of total membrane volume

^4^ – Diffusion constants based on Lommerse et al. (Lommerse, et al., 2006) and estimated from molecular weight (Andrews, 2017).

^5^ – Protein association kinetic rates based on common rates described by Schreiber et al. (Schreiber, et al., 2009)

***Data Analysis***

Data analyses were performed using R (v 3.6.1) and visualized with the ggplot2 package (v. 2.3.2.0).

**Benchmarking of particle movement and reactions**

*Particle diffusion in Cell4D accurately models Brownian motion as predicted by Fick’s Laws*

To validate the accuracy of Brownian motion for point particles within Cell4D, 100 particles were placed in the middle of a simulation space defined by 5 x 5 x 5 c-voxels, over a range of space scale and time scale parameters and allowed to diffuse. Timescales of 0.1 to 100 μs were simulated for a total of 0.1 s and the root mean squared displacement (RMSD) value of the particles were calculated at every 0.002 s interval and compared to theoretical values (**Figure 2A**). The RMSD values of the simulated particles were consistent (within 5%) with theoretical RMSD across all timescales. RMSD error showed no change over the course of a simulation, suggesting it is independent of both the number of simulation steps and overall length of the simulation. Further, we found that the accuracy of point particle diffusion was independent of simulation space scale (**Supplementary Figure 1**).

*Diffusion of bulk molecules is accurately captured by Cell4D, with simulation duration-dependent and space-scale-dependent errors*

Similar to the particle RMSD validation experiments, approximately 0.16 mM (equivalent to 100 molecules) of bulk molecules were placed in the center of a simulation space composed of 5 x 5 x 5 c-voxels of length 1µm, and allowed to diffuse outwards for 0.1 seconds. The distance of every voxel from the origin was calculated every 0.002 seconds, and the concentration of molecules within each voxel along with the distances of each from the origin were used to calculate the RMSD at every time-step using the formula:

$RMSD= \sqrt{\frac{1}{n}\sum_{i=1}^{n} \left( \mathrm{Molecules}_{i}\times d_{i} \right)^{2}}$

where *i* represents a voxel within the space, *n* is the total number of c-voxels in the simulation, and *d_i_* is the distance of voxel *i* from the center origin voxel.

Different timescales were tested with this scheme to show the performance of this method with increasing time-step lengths (**Figure 2B**). At simulation time intervals of 0.1 to 100 μs seconds, the RMSD of the molecules closely match with the theoretical dispersion of Brownian particle diffusion but appeared to diverge over time resulting in approximately 5% error after 0.1 seconds. Interestingly, the bulk molecule RMSD diverges at the exact same rate across all four timescales. By analyzing the RMSD error as a function of both time as well as number of timesteps, the error has a timestep length dependency where longer timesteps produced a larger error per step. However, the RMSD produced from the accumulation of errors over the simulation remained the same regardless of the timestep length taken, thus the error scales with simulated duration rather than time step length.

Next, the space-scale of the simulation (i.e., size of c-voxels) was increased to test the robustness of the bulk molecule diffusion simulation method and to see whether space-scale exerts an effect on simulation accuracy. Voxel lengths of 2 to 20 μm were tested with an extended duration of 1 simulated second at 0.1 ms timesteps to examine long term RMSD accuracy (**Supplemental Figure 1**). While we found the smaller space-scales (2 and 5μm) yielded a higher RMSD than the theoretical Brownian diffusion values over the course of a simulation, the error was relatively low (~5%). Further, increasing the space scale to 20μm resulted in negligible error. The error associated with small c-voxel volumes, arises as a consequence of the deterministic bulk molecule diffusion model assuming that molecules within each c-voxel are located at its center. This assumption neglects the potential random position of molecules within that space. For larger c-voxel volumes, each c-voxel will generally contain more molecules whose average position will likely reflect the center of the c-voxel, thus resulting in reduced RMSD error. For smaller c-voxel volumes, the potential RMSD error increases due to the greater impact of the random positioning of a reduced number of particles.

Overall, across all conditions tested, we found that the RMSD error did not exceed 5%, and that reducing the overall simulation time or increasing the volume of voxels, effectively eliminates errors in simulated particle and bulk diffusion.

*Unimolecular reaction kinetics are accurately simulated in Cell4D*

To validate the ability of Cell4D to accurately simulate unimolecular reactions, we defined a model reaction in which a dimer (AB) composed of two molecules, A and B, spontaneously dissociates into its component molecules. We then used Cell4D to model this reaction for 500 dimers over 0.01 seconds within a simulation environment composed of 6 × 6 × 6 c-voxels of length 0.1 μm. Results from simulations were compared to theoretical counts calculated from the integrated first-order rate law. To examine the robustness of the Cell4D reaction mechanism, different dissociation rate constants as well as simulation timestep lengths were tested. For this unimolecular reaction test, reaction rate constants from 1 × 10^0^ s^-1^ to 1 × 10^4^ s^-1^ were tested at 10-fold intervals across timescales of 0.1, 1, and 10 µs for 0.01 simulated seconds (**Supplemental** **Figure 3**). For all tested reaction constants across multiple timescales, simulated product formation matched theoretical yields indicating that Cell4D accurately simulates unimolecular first-order reactions.

*Hybrid bimolecular reactions involving bulk molecules and point particles are accurately simulated in Cell4D*

To validate the ability of Cell4D to accurately simulate reactions between bulk molecules and point particles, we defined a model reaction in which bulk molecule A can associate with point particle B to form a dimer (AB). We defined a system in Cell4D in which 500 molecules of particle B were randomly placed within a simulation environment composed of 6 x 6 x 6 voxels of length 0.1 μm (total volume 2.2 × 10^-19^ m^3^). Within this environment we further define an initial concentration for A as 3.85μM (equivalent to 500 instances of molecule A). Simulations were performed for three different reactant diffusion constants (1 × 10^-11^, 1 × 10^-12^, and 1 × 10^-13^m^2^/s) with a range of rate constants varying from 1 × 10^3^ to 1 × 10^9^ M^-1^s^-1^. Furthermore we investigated the effect of three timescales (0.1, 1, and 10µs) for a total simulation time of 0.01s. Because this hybrid model complicates the interpretation of the integrated rate law for a second-order reaction, we took advantage of a special case in which: 1) both reactants have the same concentration at Δt = 0; and 2) react with the same stoichiometry, to calculate the formation of products over time to compare to the results from Cell4D. From these simulations, we found that the accuracy of simulated reaction kinetics decreased as the reaction rate increased, with a reaction rate of 1 × 10^9^ M^-1^s^-1^ resulting in a marked deviation from theoretical values (**Supplemental Figure 4**). While timescale had negligible impact, accuracy increased for simulations with higher rates of diffusion.

*Smoluchowski method for particle-based bimolecular reactions produced errors correctable using the Andrews-Bray adjustment*

While bulk molecules are simulated differently from point particle molecules, the simulated rate of bulk molecule reactions should match the reaction rate of the molecules simulated as particles. Accordingly, using the same reaction, simulation environment and parameters defined in the previous section we examined the ability of point particles in Cell4D to accurately simulate the dimerization of two molecules A and B. We found that the particle-based reactions exhibited greater deviation from theoretical values than the hybrid reaction (**Supplemental Figure 4D**). Accuracy increased for simulations incorporating the smallest timescales (0.1 µs) and fastest reaction rates (1 × 10^9^ M^-1^s^-1^). For most of the other conditions, the simulated results underpredicted theoretical values of product formation, with errors exceeding 10% RMSE. To correct these errors we implemented the Andrews-Bray (AB) adjustment of the Smoluchowski method which has previously been shown to better simulate bimolecular reaction rates in mesoscale simulations with timescales between 1 ns to 1 ms (Andrews and Bray, 2004). The AB method uses a lookup table to approximate the theoretical binding radius of a reaction based on the average diffusion step length of the reactants and the experimentally determined rate of the reaction thus taking timescale into account by producing different binding radii depending on the length of each time step in the simulation. This method delivered high accuracy predictions for most simulations across the timescales, diffusion rates, and reaction rates tested (**Supplemental Figure 5**). We note that this method does not perform well for combinations of high reaction rates and slow rates of diffusion rates. However such conditions violate the assumption that for Smoluchowski reaction systems, the main constraint for reaction rates stems from particle collision.

**Modeling Calcium:Calmodulin binding dynamics**

*Calcium microdomains play a key role in calmodulin activation*

The C- and N-termini of calmodulin, each comprise a pair of Ca^2+^ binding EF-hand domains (C-lobe and N-lobe) that are separated by a flexible linker. Binding of Ca^2+^ to all four domains results in activation of calmodulin and the regulation of CEACAM1 functionality through binding and dissociating cis-homodimers of CEACAM1 (Edlund, et al., 1996; Patel, et al., 2013). Binding of Ca^2+^ at each lobe changes the conformation of calmodulin (T-to-R transition), resulting in cooperative binding kinetics. These kinetics are thought to involve a two-step binding model in which binding at any site of a domain increases the affinity of calcium at the other site at the same lobe (Faas, et al., 2011). Interestingly, the mechanism driving cooperativity differs between the two lobes. For the N-lobe, the T-to-R transition strongly increases k_on_ and modestly decreases k_off_. In the C-lobe, k_on_ is essentially unchanged and cooperativity results from a 1/400 decrease in k_off_. Using an ODE model, it was proposed that cooperative binding at the C-lobe supports persistence of ‘primed’ calmodulin (bearing two bound Ca^2+^) which can be achieved under modestly elevated calcium concentrations in bulk solution (~1-10µM). Such priming would then increase the likelihood of binding at all four domains, enhancing the ability of calmodulin to respond to local fluctuations in [Ca^2+^].

**Supplemental figure legends**

Supplemental Figure 1: RMSD values of Cell4D particle and bulk diffusion across multiple timescales.

A) Comparison of particle RMSD values in green with theoretical Brownian RMSD in orange. Shaded areas indicate standard deviation of particle RMSD across 5 replicates. B) The percentage error between simulated and theoretical RMSD at each measured time interval. C) Same data as B) but shown as a function of simulated time length in seconds. D) Comparison of bulk molecule RMSD values in green with theoretical Brownian RMSD in orange, with RMSD remaining the same regardless of the simulation timestep length. Theoretical values were predicted by Fick’s laws to examine timestep dependent effects of the bulk diffusion algorithm. E) The percentage error between simulated and theoretical RMSD at each measured time interval, shown relative to the number of simulated timesteps. F) Same data as E) but shown as a function of simulated time.

Supplemental Figure 2: RMSD of Cell4D bulk molecule diffusion and particle diffusion across space scales.

A) Comparison of bulk molecule RMSD (blue) and particle RMSD (green) across multiple space scales with theoretical Brownian RMSD (orange). B) The RMSD error of bulk molecules as a function of simulation space scale. C) The RMSD error of particle diffusion as a function of simulation space scale.

**Supplemental Figure 3:** **Reaction products generated from unimolecular and two-particle bimolecular reactions over time.**

A) Generation of products from a unimolecular reaction as a function of time using a time step of 10µs. The dashed lines are the predicted product concentration over time, calculated from the first-order integrated rate law. The solid lines show results from Cell4D simulations, averaged over 5 replicates. Shaded areas indicate standard deviation. B) and C) as for A) but using time steps of 1µs and 0.1µs respectively. D) Generation of products from a bimolecular reaction, using the Smoluchowski method, as a function of time using a time step of 10µs. Dashed lines are the predicted number of products as calculated through mass-action kinetics. Solid lines show results from Cell4D simulations averaged over 5 replicates. Shaded areas indicate standard deviation. Line colors indicate rate constant of the reaction, with darker colors representing faster rates. Shown are results for simulations using a particle diffusion constant of 1 × 10^-13^ m^2^/s. Under these parameters, reaction rates up to 3 × 10^6^ M^-1^s^-1^ are simulated accurately, E) and F) as for D) but using time steps of 1µs and 0.1µs respectively. For faster reaction rates, the simulations fail to model the results predicted from mass-action kinetic equations.

**Supplemental Figure 4: Summary of bimolecular products over time of bulk-particle reactions using a Smoluchowski-based method to calculate reaction probability**.

Dashed lines represent the theoretical yield of products from a bimolecular reaction involving a particle reactant and a bulk molecule reactant over 0.01 seconds. Solid lines represent yield predicted in simulations using Cell4D (mean over 5 replicates). Line colors represent the rate constant of the reaction, with darker colors representing faster rates. Time step lengths of 0.1, 1, and 10 μs were used as simulation parameters, and particle diffusion constants of 1 × 10^-11^ to 1 × 10^-13^ m^2^/s were investigated. Little or no product formation is predicted for reaction rates less than 1 × 10^6^ s^-1^. Shaded regions represent the standard deviations.

**Supplemental Figure 5: Summary of two-particle bimolecular products over time using the Andrews-Bray adjusted Smoluchowski method to calculate reaction radii.**

Dashed lines represent the theoretical yield of products from a bimolecular reaction involving two particles over 0.01 seconds. Solid lines represent yield predicted in simulations using Cell4D (mean over 5 replicates). Theoretical yields were obtained from from mass action kinetics using the equation $rate=-\frac{d\left[ A \right]}{dt}=-k \times\left[ A \right]^{2}$. Simulations used the Andrews-Bray adjusted Smoluchowski method. Line colors represent the rate constant of the reaction, with darker colors representing faster rates. Time step lengths of 0.1, 1, and 10 μs were used as simulation parameters, and particle diffusion constants of 1 × 10^-11^ to 1 × 10^-13^ m^2^/s were investigated. For reactants with 1 × 10^-11^ m^2^/s diffusion rate, the simulated product curve closely aligns with the theoretical yields at all reaction rates. For diffusion rates of 1 × 10^-12^ m^2^/s, simulations are accurate for reactions up to 1 × 10^8^ M^-1^s^-1^. For diffusion rates of 1 × 10^-13^ m^2^/s, simulations perform poorly.

**Supplementary Figure 6: Saturation of calmodulin under varying calcium concentrations in well-mixed or microdomain conditions.**

A) Proportion of calmodulin saturated with Ca^2+^ ions at equilibrium, with the number suffix denoting the number of Ca^2+^ bound to the calmodulin protein. The dotted and solid lines indicate source of data (Euler-forward deterministic prediction and Cell4D simulations, respectively). Vertical lines mark the average Ca^2+^ concentration within a T cell at rest, activated, and within Ca^2+^ microdomains. Error bars represent standard deviation of Cell4D simulations over 5 replicates. Euler-forward prediction values were calculated at 0.16 µM intervals (50 particles) for a total of 101 data points. B) Comparison of CaM saturation in volumes under well-mixed or microdomain conditions at 0.81, 0.97, 1.14, 1.3, 1.46, and 1.62 µM, with the dotted, dashed, and solid lines representing the predicted well-mixed equilibrium, Cell4D simulated well-mixed results, and Cell4D simulated spatial microdomain conditions. CaM_2 (in yellow) shows the proportion of calmodulin with two bound calcium binding sites, and CaM_4 (green) represents fully saturated calmodulin. Error bars represent standard deviations of Cell4D simulations over 5 replicates.

Supplementary Figure 7: Comparing surface CEACAM1 clustering in both Cell4D model variants.

The amount of clustered membrane monomeric CEACAM1 over 10 simulated seconds is compared between the raft and no-raft models across several calmodulin and Lck concentrations. No clustered CEACAM1 can be found in the calmodulin-absent systems, and the concentration of clustered molecules is positively correlated with calmodulin concentration. In the absence of Lck, the raft model produced more clustered proteins while the opposite is true when Lck is present.
